# *Trp53* heterozygosity defines novel states impacting tumor plasticity and progression in Kras-initiated non-small cell lung cancer

**DOI:** 10.64898/2026.10.12.758643

**Authors:** V Mitheera, Abhishek Koladiya, Aatish Thennavan, Xiaojie Yu, Guillermina Lozano

## Abstract

*KRAS* and *TP53* are frequently mutated in non-small cell lung cancer (NSCLC), the leading cause of cancer-related mortality worldwide. The most common event at the *TP53* locus is mutation/loss of one allele and retention of a wild-type allele, which confers better survival outcomes than complete *TP53* loss. To study the contribution of *TP53* heterozygosity, we generated a murine lung tumor model with a *KrasG12D* mutation and intact (*K*) or single-allele loss (*KP+/*-) of *Trp53 and* compared it with publicly available complete loss (*KP*) data. K and KP+/− mice lived longer than KP mice, paralleling clinical survival data. Single-cell transcriptomic analysis of tumor-burdened lungs indicated that *Trp53* heterozygosity expanded transcriptional plasticity that was not present in K and KP tumors. Genomic instability was also higher in KP+/− mice relative to the KP genotype. *Trp53* gene dosage, thus, regulates phenotypic plasticity and evolutionary trajectories of tumor states across K, KP+/−, and KP tumors. Murine KP+/− tumor signatures and copy-number profiles were present in human NSCLC datasets. Two of these signatures correlated with better and worse patient survival outcomes, respectively. *TP53* heterozygosity in *KRAS*-mutant lung cancers presents a distinct genomic landscape that should be taken into account in the development of novel therapeutic strategies.

## Introduction

Lung cancer is the leading cause of cancer-related mortality globally, accounting for nearly one in five cancer deaths annually^1^. Lung cancers are broadly classified into two major categories – small cell lung cancer (SCLC) and non-small cell lung cancer^2^ (NSCLC). NSCLC accounts for 80-85% of all lung cancers. NSCLC can be subdivided into three main subtypes: lung adenocarcinomas (LUAD), lung squamous cell carcinomas (LUSC), and large cell carcinomas^2^(LCC). The most frequently altered genes in NSCLC are *TP53*, followed by *KRAS* and *EGFR*, with enrichment of *KRAS* or *EGFR* activating mutations in LUAD^3–5^. Despite advances in targeted therapies for *KRAS*-mutant and *EGFR*-mutant LUAD, disease recurrence rates remain high, likely caused by acquired resistance. This is particularly enriched in patients treated with KRAS inhibitors^6–8^.

Co-occurrence of *KRAS* and *TP53* mutations in patients significantly reduces median overall survival, reflecting cooperativity in driving aggressive LUAD^9,10^. *TP53-KRAS* co-mutant LUADs are “immune-hot” with an inflamed tumor microenvironment, increased tumoral PD-L1 expression, and higher tumor mutational burden in patients^11–13^. Hence, patients with advanced KRAS/TP53-mutant LUAD may receive immune checkpoint inhibition (ICI) as part of standard systemic treatment^14,15^. However, LUAD patient response to ICI correlates with the expression of tumoral PD-L1 only if more than 50% of tumor cells express the ligand^16^. Hence, there is an unmet need to elucidate the progression of *KRAS-TP53* co-mutant LUAD to unveil more targeted therapies.

To examine the cooperativity of *KRAS* and *TP53* mutations in driving aggressive forms of LUAD, genetically engineered mouse models (GEMMs) with an activating *Kras^G12D^* mutation and *Trp53* deletion (*Kras^G12D/+^*;*Trp53^flox/flox^*[KP]) were created to model lung cancer progression and evolution *in vivo*^17^. The KP model has faithfully recapitulated the clinicopathological progression of LUAD from adenomas to adenocarcinomas. Further studies utilizing the KP model (and other functionally comparable GEMMs) have identified the role of wild-type p53 in limiting the progression of lung adenomas to adenocarcinomas and defined the near-stereotypic progression of lung tumors^18–21^.

While the KP model has been invaluable in revealing the diverse tumor states arising from complete loss of *Trp53* in Kras-initiated LUAD, it only assays for p53 loss-of-heterozygosity (LOH). In human LUAD, *TP53* LOH occurs in advanced disease^22^ but is rare in most progressing lesions^23^ (less than 1%). Indeed, monoallelic loss of *TP53* (heterozygosity) is a more common genetic alteration in these patients. Thus, the KP model bypasses the *TP53* heterozygous state commonly observed in NSCLC patients. Furthermore, the timing of *TP53* LOH (early or late in tumor progression) and its consequences for tumor evolution remain unknown. Therefore, a critical knowledge gap exists in how progressive allelic loss of *TP53* regulates tumor progression and evolution in LUAD.

To determine if there are differences in the progression of Kras-initiated lung tumors arising in *Trp53*-heterozygous mice in comparison to mice developing LUAD with complete loss of *Trp53* (KP) or with intact wild-type *Trp53* (K), we generated the *Kras^G12D/+^*; *Trp53^flox/+^* heterozygous (KP+/−) mouse model. Intratracheal injection of lentivirus expressing Cre recombinase in an adult mouse lung led to sporadic and somatic induction of oncogenic Kras^G12D^ mutation and varying doses of p53.

Single-cell RNA (scRNA) sequencing of tumor-bearing K and KP+/− mice lungs and comparison to publicly available murine KP tumor scRNA datasets identified unique cellular states arising in *Kras*-mutant tumors initiated with differing doses of *Trp53*, with KP+/− tumors reflecting a divergent tumor evolutionary path through the acquisition of novel states defined by copy-number variations and phenotypical plasticity. In addition, tumor progression in KP+/− mice paralleled that observed in LUAD patient data. Furthermore, select gene signatures derived from the KP+/− model held prognostic power in stratifying survival outcomes in human *KRAS*-mutant LUAD clinical data, highlighting the translational relevance of modeling Kras-mutant lung tumors with clinically relevant *p53* dosage *in vivo*.

## Results

### p53 dosage determines survival and pathological progression in Kras-mutant lung cancers

To investigate the role of *TP53* allelic gene dosage in the progression of *KRAS*-mutant human NSCLC, we first analyzed clinical data from patients with *KRAS*-mutant NSCLC and stratified them according to *TP53* gene dosage^24,25^. Patients harboring wild-type *TP53* had the longest overall survival (n = 1500; 54.3 months), followed by patients with single-allele TP53 loss (n = 100; 35.6 months), whereas patients with complete *TP53* deletion had the shortest median overall survival (n = 4; 18.3 months) (Supp Fig 1A). The observed decrease in median survival from 54.3 months in TP53-wild-type tumors to 35.6 months in tumors with monoallelic *TP53* loss supports a clinically relevant association between reduced *TP53* dosage and poorer outcome. Moreover, monoallelic *TP53* loss was substantially more frequent than complete *TP53* deletion among KRAS-mutant NSCLC patients. Since only four *KRAS*-mutant NSCLC patients showed a complete loss of *TP53*, statistical comparisons involving this group were underpowered. Together, these findings highlight the clinical relevance of the *TP53*-heterozygous state and provide a rationale for investigating how monoallelic *TP53* loss influences the progression and evolutionary trajectories of KRAS-mutant lung tumors.

To directly determine whether the phenotypic consequences of *TP53* allelic gene loss in human KRAS-mutant NSCLC could be recapitulated in spontaneous murine models of lung cancers, we generated cohorts of *LSL-Kras^G12D/+^* (n = 14) and *LSL- Kras^G12D/+^;Trp53^Flox/+^* (n = 14) genetically engineered mouse models (GEMMs). Intratracheal administration of a lentivirus expressing the *Cre* recombinase^17^ (Fig 1A; Supp Fig 1B) initiates lung tumors by activation of oncogenic *Kras^G12D/+^* mutation alone (subsequently called K mice) and concomitant *Kras^G12D/+^* mutation and mono-allelic deletion of *Trp53* (labeled KP+/− mice). Tumor-free survival was calculated for moribund K and KP+/− mice succumbing to lung disease, revealing 1) a vastly greater tumor latency and tumor-free survival times in K and KP+/− mice in comparison to KP^17^ (endpoint of 6 months), and 2) 60% of K mice were alive at 20 months when all KP mice had died (Supp Fig 1C). We then isolated tumor-bearing lungs from K and KP+/− mice and performed lung tumor pathology to evaluate whether *Tr*p53 dosage influenced tumor burden and grade. Multifocal lung tumors were observed in moribund K and KP+/− mice. (Supp Fig 1D). Gross morphological examination of tumor-bearing K and KP+/− lungs revealed differences between *Trp53* gene dosage and tumor burden: K lungs exhibited discrete, multifocal nodules, whereas KP+/− lungs showed macroscopic tumor coalescence and significantly larger primary lesions (all mice) (Supp Fig 1D). Histopathological analysis of K and KP+/− tumor-bearing lungs supported this finding, with predominantly adenomas in K (n = 1) tumor-bearing lungs and adenocarcinomas present only in KP+/− (n = 2) lungs (Supp Fig 1E). Quantification of tumor burden in K and KP+/− tumor-bearing murine lungs revealed almost a 4-fold higher tumor burden in KP+/− compared to K lung (Supp Fig 1F). Thus, both tumor grade and volume were significantly higher in *Kras*-mutant mice with *Trp53* heterozygosity than in *Kras* mice.

**Fig 1:**
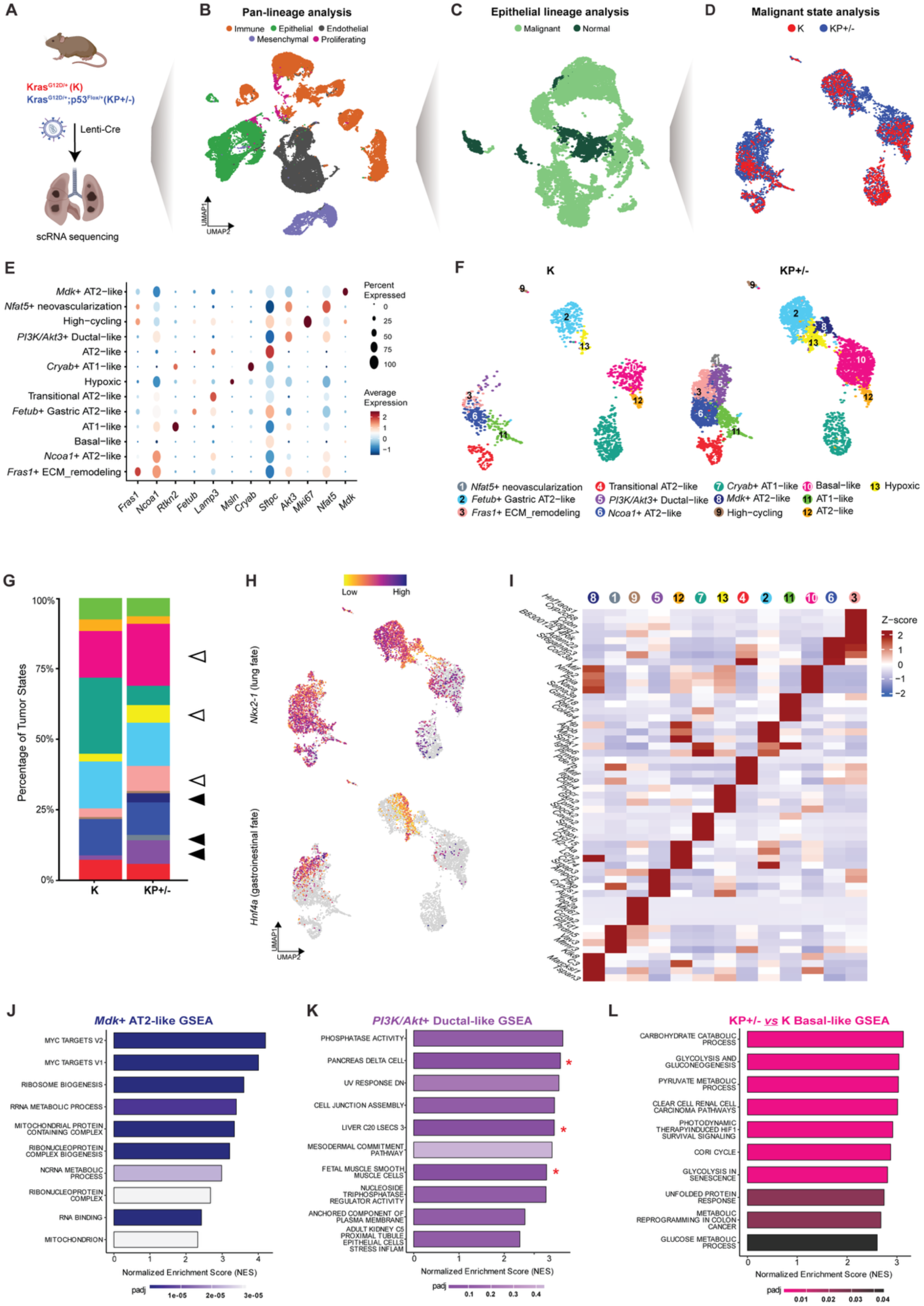
Single-cell profiling of K and KP+/− tumors revealed distinct cell states. **(A)** Schematic diagram of experimental strategy to initiate lung tumors through intratracheal instillation of lentiviral Cre in *Kras^G12D/+^* (K) and *Kras^G12D/+^*; *p53^Flox/+^* (KP+/−) mice. Lungs were collected from moribund mice for pathological and single-cell analyses. **(B)** The first step in the single-cell analyses workflow identifies major cell lineages in scRNA-sequencing data of K and KP+/− tumor- bearing lungs: Epithelial (green), Endothelial (charcoal), Immune (orange), Mesenchymal (purple), and Proliferating (magenta). **(C)** The second step in this workflow distinguishes malignant (light green) from normal (dark green) epithelial cells. **(D)** The final step involves only malignant cells in K (red) compared to KP+/− (blue) tumors. **(E)** STRING pathway interaction network analysis reveals developmental and metabolic differences, and perturbations in *PI3K/Akt* receptor-growth factor signaling in KP+/− malignant cells compared with K counterparts. **(F)** High-resolution annotation of KP+/− and K malignant cells unveils thirteen distinct states. **(G)** The stacked bar plots show the percentage of each tumor state comprising K and KP+/− malignant cells, where filled black arrowheads identify unique states exclusive to KP+/− tumors, and empty arrowheads indicate K states that expanded in KP+/− tumors. **(H)** UMAP visualizations of cells reflect the expression of *Nkx2-1* (lung fate marker) and *Hnf4a* (primordial gut fate marker). **(I)** Heatmap reveals the Z-scores of the top 4 differentially expressed genes defining each of the 13 tumor states from (F). **(J-K)** Normalized Enrichment Scores (NES) from Gene set enrichment analyses (GSEA) identify functional pathways enriched in states exclusive to KP+/− tumors: *Mdk*+ AT2-like (J) and *PI3K/Akt*+ Ductal-like (K). Statistically significant enrichment of pathways in (K) is marked by a red asterisk. **(L)** Normalized Enrichment Scores (NES) from Gene set enrichment analyses (GSEA) identify functional pathways enriched in the Basal-like tumor state that expanded in KP+/− tumors compared to K. and the unfolded protein response (UPR) (Fig 1L, Supp Fig 4E, Supp Table 4). As for the KP+/− exclusive *Nfat5*+ neovascularization state, most of the enriched neuronal pathways were not statistically significant, possibly due to the small number of cells (n = 89) in this cluster, but the leading genes overrepresented in these pathways were expressed predominantly in this cluster (Supp Fig 4F, G, Supp Table 3). In the Hypoxic and *Fras1*+ ECM_remodeling states, hypoxia and basal cell carcinoma were the top pathways enriched with corresponding hypoxic (*Ak4*) and ECM engagement (*Eya4*) genes upregulated in the KP+/− compared to the K counterparts (Supp Fig 4H-K, Supp Table 4). Collectively, the characterization of KP+/− states suggests that discrete groups of malignant cells emerge, each utilizing distinct signaling pathways and developmental programs to potentiate intertumoral transcriptional heterogeneity.

### Identification of malignant cells in single-cell transcriptomic profiling of K and KP+/− tumor-bearing lungs

To elucidate the full spectrum of transcriptional states arising in *Kras*-mutant tumors initiating with wildtype or single-allele loss of *Trp53 in vivo*, we collected tumor-bearing K (n = 3) and KP+/− (n = 3) lungs from moribund mice for single-cell RNA (scRNA) sequencing (Fig 1A; Supp Fig 1B). A total of 21361 and 22819 single cells were sequenced from K and KP+/− tumor-bearing lungs, respectively (Fig 1B, Supp Fig 2A, Supp Table 1). Single-cell datasets were integrated using Harmony for downstream analyses^26^. To identify the major cell lineages in K and KP+/− tumor-bearing lungs, we performed lineage analyses guided by the expression of canonical lineage markers: *Plvap*, *Icam2,* and *Cdh5* for endothelial, *Ptprc* for immune, *Cdh1*, *Epcam*, *Krt8,* and *Krt18* for epithelial, and *Col1a1*, *Col3a1*, *Pdgfra* and *Pdgfrb* for mesenchymal lineages. *Mki67* identified proliferating cells (Supp Fig 2B). The percentage of cells comprising the five lineages revealed an expansion of the epithelial compartment in KP+/− tumor-bearing lungs compared to K counterparts (32.2% vs. 13.9%) (Supp Fig 2C). In addition, the KP+/− immune compartment was reduced in proportion to the K counterpart (46.9% vs. 30.0%). Next, we subclustered the K and KP+/− epithelial lineage, since pathological analyses confirmed that tumors in these mice (adenomas and adenocarcinomas) were of epithelial nature. High-resolution clustering generated 26 clusters (Supp Fig 3A). To systematically differentiate the clusters representing normal epithelial cell types from malignant populations, we utilized the LungMAP murine reference database for automated annotation^27^ (Supp Fig 3B). This analysis mapped epithelial clusters 2, 17, and 21 as alveolar type 1 (AT1) cells, clusters 3, 7, 8, 15, 16, and 19 as alveolar type 2 (AT2) cells, clusters 13 and 25 as airway ciliated cells, and cluster 18 as airway club or secretory cells (Supp Fig 3A - B). To ensure high-confidence identification of normal epithelial cells in our data, we calculated enrichment scores of clusters from previously published epithelial cell signatures of AT2^28^, AT1^28^, Secretory^29^ (airway club), and Ciliated^29^ cells (Supp Fig 3C- F). This approach enabled us to identify normal epithelial cell types based on the highest--scoring cluster for each of the four signatures. The remaining epithelial clusters lacked a consistent normal epithelial identity across these complementary annotation approaches and were therefore operationally classified as malignant tumor-associated epithelial populations for downstream analyses (Fig 1C, Supp Fig 3G, Supp Table 1). We further validated this classification by evaluating the expression of canonical genes defining the major normal cell types in the K and KP+/− epithelial compartments, demonstrating that these markers were expressed only in the four normal epithelial cell types (Supp Fig 3H).

### Trp53 gene dosage impacts tumor transcriptional heterogeneity

To identify the biological pathways overrepresented in KP+/− compared with K malignant cells, we performed STRING pathway and process enrichment analyses using differentially upregulated genes (DUGs) that were at least 2.5-fold significantly increased in expression in KP+/− compared with K malignant cells^30^ (Supp Fig 4A). The resulting network map revealed clusters with nodes or pathways related to 1) receptor-growth factor and integrin signaling, including *PI3K*/*Akt* signaling, 2) pancreas and endodermal development, and 3) epithelial structure development and new blood vessels (Fig 1E). Other distinct, functional clusters include various metabolic pathways – including those of glucose, amino acids, soluble lipids, and vitamins (Supp Table 2). Kidney-related pathways and neural cellular projections were also evident and are commonly observed in murine lung cancers undergoing lineage infidelity^21^. Thus, the most enriched Gene Ontology pathways in KP+/− tumors were developmental processes, thereby facilitating lineage infidelity (Supp Fig 4B).

Clustering of K and KP+/− malignant cells yielded thirteen discrete states (Fig 1F). Tumor states were annotated based on the expression of human NSCLC prognostic genes (*Nfat5*, *Fetub*, *Fras1*, *Akt3*, *Ncoa1*, *Mdk*)^31–36^ and published signatures of lung alveolar and gastrointestinal cell types, cell cycle, and hypoxia (Fig 1E, Supp Fig 4C)^37–39^. Tumor states also exhibit differences in transcriptional breadth, as measured by the number of genes detected per cell across states (Supp Fig 4D). Analysis of tumor states in K and KP+/− malignant cells identified *Mdk*+ AT2-like (cluster 8), *Nfat5*+ neovascularization (cluster 1), and PI3K/Akt Ductal-like (cluster 5) as novel states exclusive to KP+/− (Fig 1F, G). In addition, the Basal-like (cluster 10), Hypoxic (cluster 13), and *Fras1*+ extracellular matrix (ECM) remodeling (cluster 3) states expanded in the KP+/− malignant compartment compared to K (Fig 1G). The expression of *Nkx2-1*, a lung epithelial lineage-defining marker, was reduced in clusters exclusive to KP/- and corresponded with the induction of regulator of gastrointestinal fate, *Hnf4a* (Fig 1H), again reflecting lineage infidelity in KP+/− tumors^21^. Finally, distinct gene markers defined the thirteen annotated tumor states (Fig 1I).

To further characterize the functional programs associated with tumor states unique to or expanding in KP+/− tumors, we performed Gene Set Enrichment Analysis (GSEA) on state-specific DUGs^40^ (Supp Table 3). Analysis of the *Mdk*+ AT2-like state (cluster 8) revealed a significant enrichment of *Myc* targets and various anabolic processes (Fig 1J), suggesting that these cells represent a *Myc*-driven, highly proliferative population. In contrast, the *PI3K/Akt*+ Ductal-like state (cluster 5) trended towards an enrichment of phosphatase activity and cell junction assembly pathways, alongside significantly upregulated gene expression patterns for pancreas delta cells and growth-factor secreting liver stellate cells (Fig 1K; red asterisks). Furthermore, KP+/− Basal-like state (cluster 10) revealed a significantly drastic metabolic shift in comparison to the K Basal-like state, as evident in the differentially upregulated metabolic genes (*Ldha*, *Pgk1*, *Pkm*, *Pgam1*, *Prmt1*) and a marked induction of carbohydrate catabolic processes, glycolysis,

### Bidirectional label transfer reveals divergent paths of progression in K and KP+/− tumors in comparison to published KP model

To systematically compare the transcriptional landscapes of K and KP+/− lung tumors against KP counterparts that lack both copies of *Trp53*, we performed label transfer using a previously published KP lung tumor dataset as a reference atlas^41^. By projecting K and KP+/− malignant cells onto the KP malignant reference annotations, we observed that K and KP+/− cells primarily mapped to AT1-like, AT2-like, and the various gastric-like states (Fig 2A, Supp Fig 5A). Strikingly, the pre-EMT, early EMT-2, and Mesenchymal-like states comprising the KP transcriptional landscape were not present in K and KP +/− malignant tumors, suggesting that K and KP+/− do not complete the EMT process (Fig 2B, Supp Fig 5B). The absence of gross metastasis in K and KP+/− lung tumor models may explain the lack of K and KP+/− malignant cell mapping to the mesenchymal-like cells isolated from metastatic lung tumors (Mesenchymal-1 (metastatic), Mesenchymal-2 (metastatic)) in KP tumors (Fig 2B). Gene expression analysis of K and KP+/− tumor state signatures across KP tumor states validated the label transfer, highlighting restricted expression of K and KP+/− signatures to the lung-like (AT1- like, AT2-like), gastric, endodermal, and high plasticity cell states (HPCS) in the KP dataset (Fig 2C, Supp Table 5).

**Fig 2:**
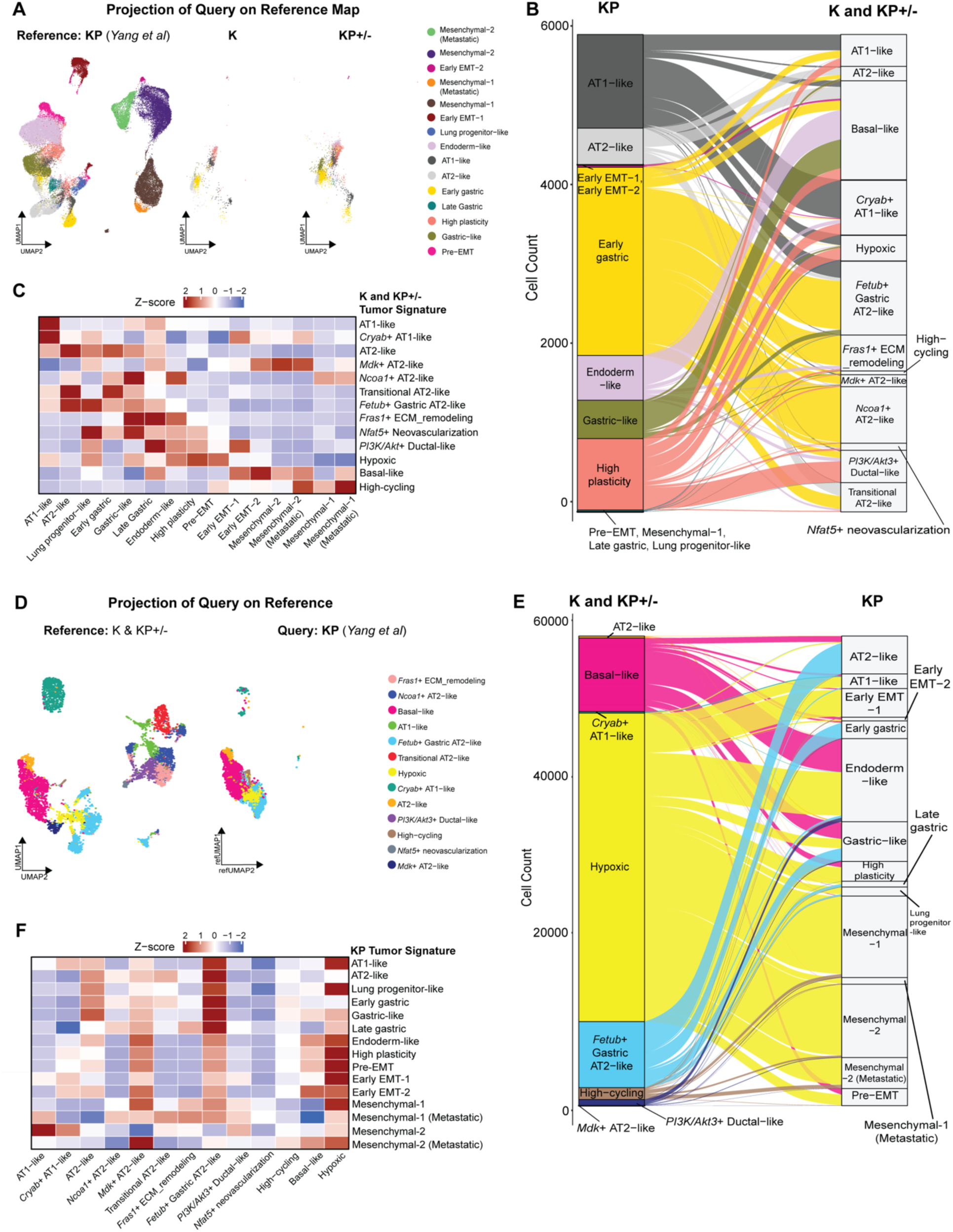
Transfer label learning identified tumor cell states unique to KP+/− and KP lung tumors. (A) UMAP projection of K and KP+/− malignant cells onto reference KP (*Kras^G12D/+^* lung tumor cells with complete loss of p53; Yang et al) dataset. Malignant cells are colored by their predicted tumor labels from the KP reference. (B) Sankey diagram maps KP reference states (colored; left) to the K and KP+/− query malignant cells (right). The flow thickness represents cell count. (C) Heatmap reflects Z-scored expression of K and KP+/− gene signatures (x-axis) across the KP tumor states (y-axis). (D) UMAP projection of KP malignant cells (Yang et al) onto reference K and KP+/− states. Query cells are colored by their predicted tumor labels from K and KP+/− reference (from Fig 1). (E) Sankey diagram depicts the transfer of labels from the K and KP+/− reference (colored; left) to the query KP malignant cells (right). (F) Heatmap of Z-scored KP tumor signature expression in K and KP+/− reference states.

Projecting KP cells onto our K and KP+/− reference malignant states, further revealed genotype-specific differences in Kras-mutant lung tumors. Firstly, KP malignant cells exhibited a high degree of transcriptional overlap with the Hypoxic, *Fetub*+ Gastric AT2- like and Basal-like clusters in the K-KP+/− tumors (Fig 2D). Second, KP tumors showed limited occupancy within the 1) novel states of KP+/− tumors (*PI3K/Akt* Ductal-like, *Mdk*+ AT2-like, *Nfat5*+ neovascularization) and other states in K expanding in KP+/− (*Ncoa1*+ AT2-like, *Fras1*+ ECM_remodeling) (Supp Fig 5C-D). The Hypoxic state in K and KP+/− tumors mapped to numerous states in the KP dataset, ranging from pre-EMT, Mesenchymal-like, Endoderm-like, Gastric and HPCS (Fig 2E). On the other hand, the *Fetub*+ Gastric AT2-like, as the name suggests, maps to the AT2-like and Gastric states in the KP dataset. Therefore, the mapping of K and KP+/− tumor states to KP tumors was less discriminative, with multiple transcriptionally distinct KP states converging onto a limited number of K and KP+/− states. This likely reflects inherent differences in the state spaces of the K – KP+/− and KP models, as each contains tumor states without direct counterparts in the other, thereby limiting one-to-one correspondence by label transfer.

Validation of this mapping was performed through a gene expression analysis of KP tumor state signatures in K-KP+/− tumor states: lung-like and gastric-like signatures mapped to the *Fetub*+ Gastric AT2-like tumor state, and the remaining (endodermal, HPCS, EMT, and mesenchymal-like) states mapped to the Hypoxic state (Fig 2F). The asymmetry observed in bidirectional label transfer further indicates that the transcriptional state spaces of these models are not equivalent, suggesting that K and KP+/− tumors do not simply represent earlier stages of the canonical KP trajectory. Therefore, our results suggest that *Kras*-mutant lung tumors that initiate with single-allele loss of *Trp53* follow a unique path of tumor progression, characterized by novel transcriptional programs that diverge from the previously elucidated progression in the KP model^21,41^.

### Genomic instability and Trp53 loss-of-heterozygosity define distinct states in K and KP+/− tumors

Next, we investigated the relationship between genomic stability and the novel transcriptional states observed in KP+/− tumors. To do so, we inferred copy number variation (CNV) across K, KP+/−, and KP genotypes, using normal T-cells as a diploid reference baseline^42^ (Fig 3A). CNV analyses revealed profoundly different genomic landscapes in K and KP+/− tumors compared to the KP counterparts. Consistent with previous studies, KP tumors displayed some genomic instability, with focal CNVs detected^21,41^ (Fig 3D). In stark contrast, both K and KP+/− tumors exhibited genome-wide chromosomal aberrations, including numerous segmental and whole-chromosome gains and losses (Fig. 3B, C).

**Fig 3:**
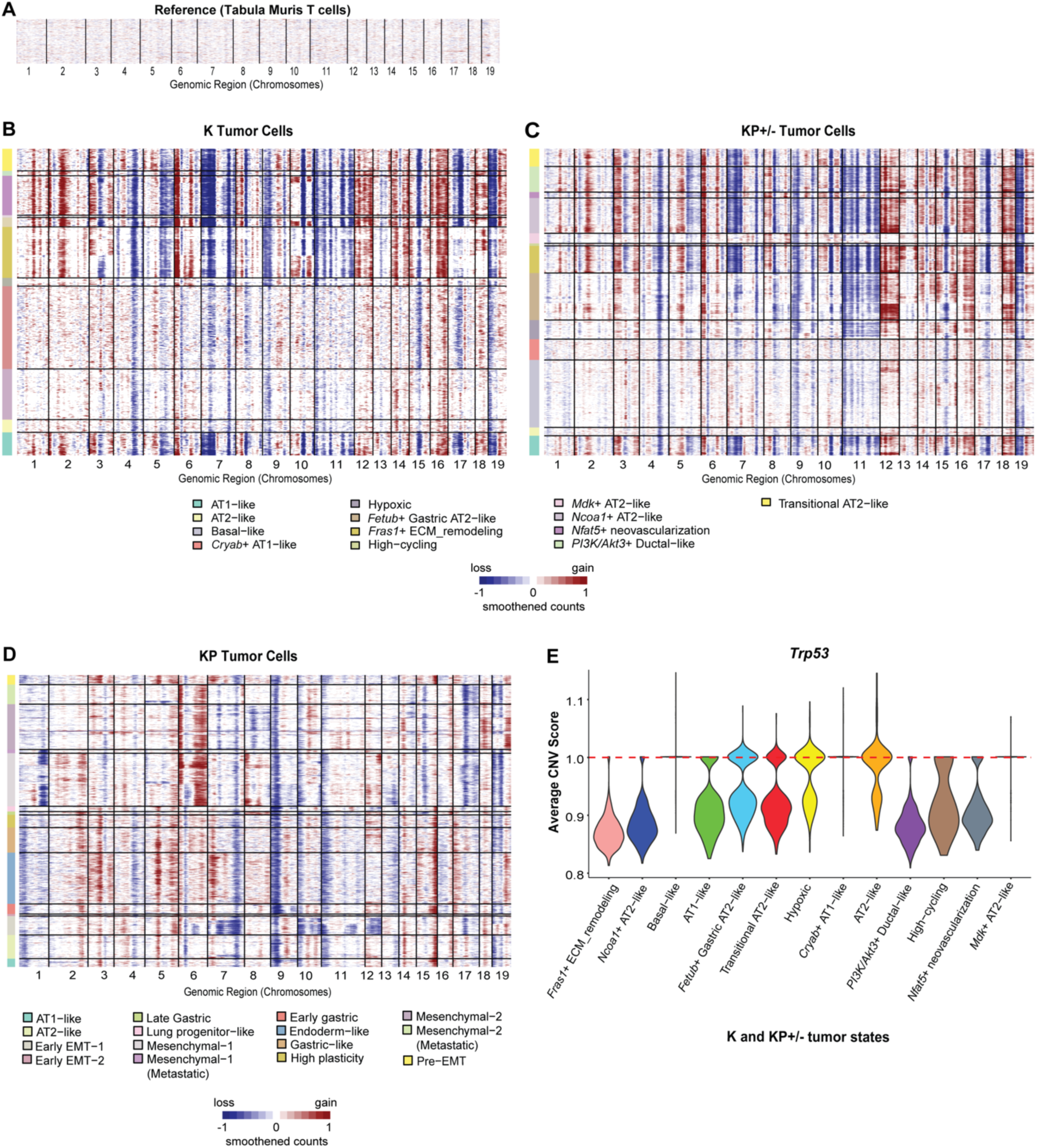
Genomic instability defines K and KP+/− tumors. **(A)** Reference map using T cells from the *Tabula Muris* atlas as a diploid baseline indicates a stable genomic profile across all chromosomes. **(B–D)** Heatmaps of inferred Copy Number Variations (CNVs) for (B) K, (C) KP+/−, and (D) KP tumor cells (*Yang et al*). Malignant cells (rows) are grouped by their annotated state (left color bar) and ordered by positions along the chromosomes (columns). **(E)** Violin plots indicate the inferred CNV score at the *Trp53* locus of malignant cells across K and KP+/− tumor states. The dashed red line at *1.0* indicates copy number neutrality.

Subsequently, we quantified this genome-wide chromosomal instability across malignant cells comprising every K, KP+/, and KP tumor state using an aggregated CNV burden score (Supp Fig 6A-C, Methods). CNV burden was not observed across all novel KP+/− and expanding states from K but only in the *PI3K/Akt*+ Ductal-like, *Nfat5*+ neovascularization and *Fras1*+ ECM_remodeling cell states, respectively (Supp Fig 6A-B). However, the tumor states shared by K and KP+/− malignant populations (AT1-like, *Cryab*+ AT1-like, AT2-like, Transitional AT2-like, *Fetub*+ Gastric AT2-like, *Ncoa1*+ AT2- like) possessed comparable CNV burden across both genotypes. Subsequently, the KP malignant states have very low CNV burden, apart from CNV events occurring in the mesenchymal-like and metastatic states (Supp Fig 6C), supporting the previous finding that the complete genetic ablation of *Trp53* in the KP model enables progression to tumor states without requiring extensive chromosomal rearrangement^41,43^. Interestingly, the states predominantly mapped from K and KP+/− malignant cells onto KP tumors (Basal-like, Hypoxic, *Fetub*+ Gastric AT2-like) also possessed lower CNV burden. Together, these results indicate that increased chromosomal instability characterizes a subset of the distinct tumor states emerging in KP+/− tumors, contributing to the expanded transcriptional heterogeneity.

Since genomic instability differentiates novel and expanding states arising in KP+/− tumors, we hypothesized that this instability might be facilitated by the stochastic loss of the remaining wild-type *Trp53* allele. To test this hypothesis, we analyzed the average inferred CNV score for all cells in the thirteen K and KP+/− clusters at the *Trp53* locus. Strikingly, almost all malignant cells in tumor clusters with the highest CNV burden (*PI3K/Akt*+ Ductal-like, *Nfat5*+ neovascularization, *Fras1*+ ECM_remodeling, *Ncoa1*+ AT2-like, AT1-like) and the *Fetub*+ Gastric AT2-like state were characterized by a marked decrease of *Trp53* CNV score of 0.85–0.95 from copy number neutrality (CNV score of 1.0) (Fig 3E). Statistical analyses confirmed that in tumor states with the highest CNV burden, *Trp53* dosage reduction was significant in KP+/− tumors compared with K counterparts (Supp Fig. 6D). We then extended this analysis to the other genetic driver in this lung tumor model, *Kras*. Although the K, KP+/−, and KP models utilize a conditionally active *Kras-G12D* mutation, we observed copy-number gains at the *Kras* locus in malignant cells across K, KP+/−, and KP states. In the K and KP+/− tumors, states with *Trp53* copy-number loss positively correlated with copy-number gains of Kras (a score of 1.1 or above) (Supp Fig 6E). In the KP model, only mesenchymal-like and mesenchymal metastatic states with detectable CNV burden acquired copy number gains at the *Kras* locus (Supp Fig 6F). Therefore, *Kras* copy number gains could result from *Trp53* LOH in the K and KP+/− tumor states and from genomic instability in the KP tumor states.

### Trp53 gene dosage regulates phenotypic plasticity and evolutionary trajectories in Kras-mutant lung tumors

To trace the evolution of K, KP+/−, and KP tumors, we first inferred the developmental stemness and plasticity of tumor states using CytoTRACE^44^ (Methods). Higher CytoTRACE scores indicate less differentiated cellular states with greater developmental potential. Tumor states novel to or expanded in KP+/− tumors (*PI3K/Akt*+ Ductal-like, *Nfat5*+ neovascularization, *Mdk*+ AT2-like, *Fras1*+ ECM_remodeling, Hypoxic, Basal-like)) exhibited a broad range of CytoTRACE scores. Several states with high inferred CNV burden, including the *PI3K/Akt*+ Ductal-like, *Nfat5*+ neovascularization, *Fras1*+ ECM_remodeling, and *Ncoa1*+ AT2-like, displayed relatively high CytoTRACE scores, with KP+/− *Fras1*+ ECM_remodeling state having a significantly higher score compared to the K counterpart (Fig 4A). In contrast, the Basal-like state has the lowest CytoTRACE score, suggesting it is one of the earlier, more differentiated states that arise during tumor progression. The CytoTRACE score for the *Mdk*+ AT2-like state falls between that of the high-CNV states and the low-CNV Basal-like state, suggesting its potential as a transitional bridge or intermediary state in the progression of KP+/− tumors. In KP tumors, the states unique to this genotype had the highest CytoTRACE scores (state names in bolded green text). These findings reflect the role of complete loss of *Trp53* in driving the progression of Kras-mutant lung tumors by giving rise to highly aggressive, transdifferentiated mesenchymal-like states that drive metastasis in the KP model^41^ (Fig 4B).

**Fig 4:**
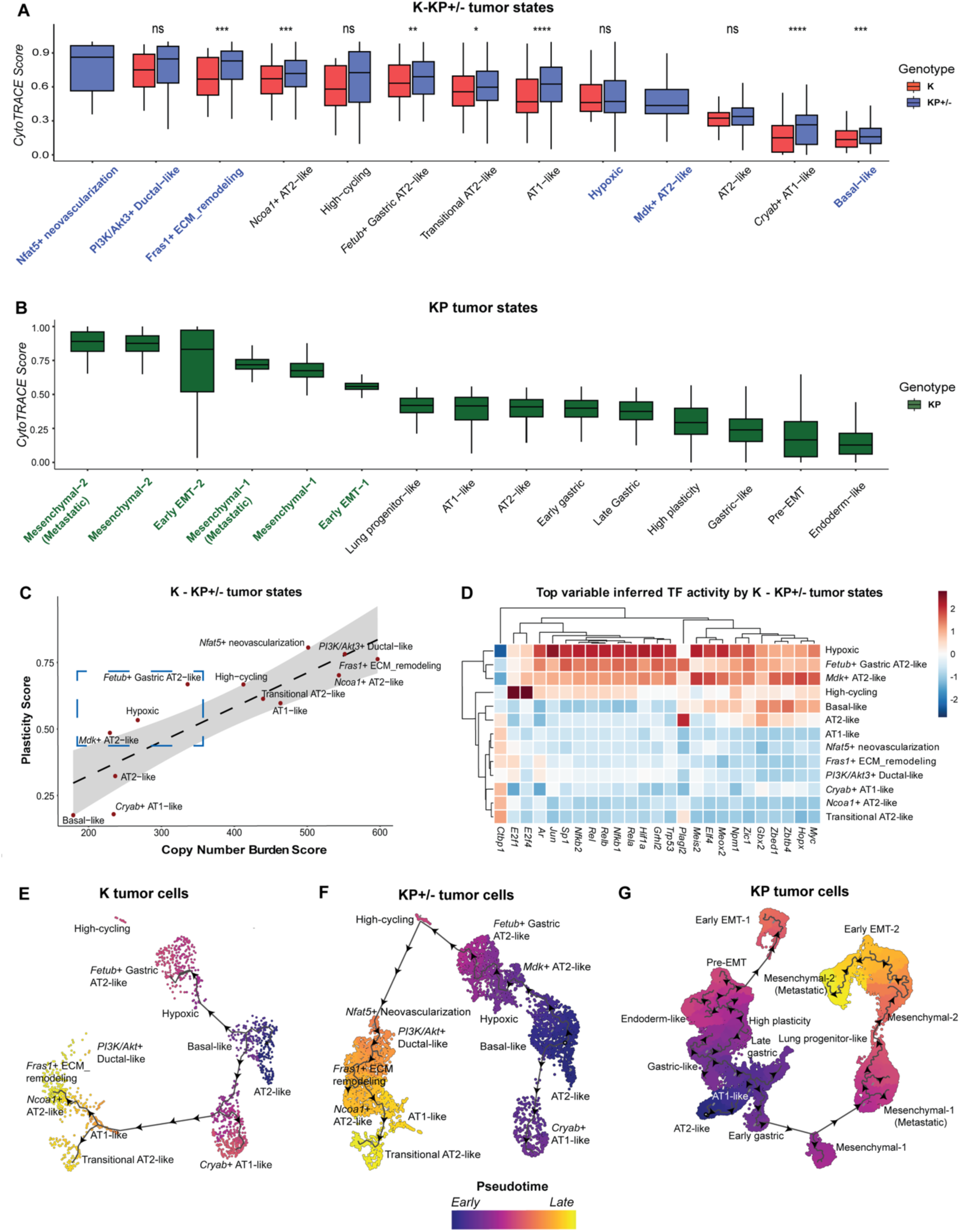
Phenotypic plasticity and evolutionary trajectories in *Kras*-mutant lung tumors vary with *Trp53* gene dosage. **(A-B)** CytoTRACE-based scoring for plasticity in (A) K and KP+/− cellular states and (B) KP counterparts. A score of 1 indicates the least differentiated and most plastic tumor state, and a score of 0 indicates a terminally differentiated cell state with the lowest plasticity potential. Bolded state names in (A) reflect states exclusive to KP+/− tumors or expanding from K tumors, and in (B) reflect states exclusive to KP tumors. **(C)** Scatterplot reflects the positive correlation between plasticity score (from cytoTRACE) and copy number burden score (from inferCNV) in K and KP+/− tumor cells. Clusters on the dotted line show a perfect linear correlation between plasticity and copy-number variation. **(D)** Transcription Factor Activity analysis reveals Z-scored scaled activity of transcription factors (TFs) that govern the K and KP+/− tumor states. **(E-G)** *Left to right*: Monocle-based pseudotime analysis depicting the evolutionary trajectories undertaken by tumor cells in (E) K, (F) KP+/−, and (G) KP tumors when the cell-of-origin is set to be AT2-like cells. The purple shade reflects tumor clusters that arise earlier in tumor evolution, and the yellow shade reflects newer states that arise later in this process.

A correlation analysis between the inferCNV and CytoTRACE scores reiterated the positive correlation between CNV burden and plasticity in KP+/− tumor states, indicating that genomic instability either drives or stabilizes phenotypic plasticity in KP+/− tumors (Fig 4C). Strikingly, the low-CNV burden KP+/− states that 1) mapped to the KP malignant cells (*Fetub*+ Gastric AT2-like, EMT) and 2) are novel to the KP+/− genotype (*Mdk*+ AT2- like) had higher plasticity relative to their CNV burden (Fig 4C; dashed blue box). Therefore, to determine whether the disproportionately elevated plasticity evident in these three states was underscored by dysregulated transcriptional networks, we inferred transcription factor (TF) activity across all 13 KP+/− tumor states (Supp Table 7). The three tumor states with the greatest inferred TF activity were also the EMT, *Fetub*+ Gastric AT2-like, and *Mdk*+ AT2-like states (Fig 4D). In particular, the *Mdk*+ AT2-like state had the highest *Myc* expression and inferred transcription factor activity (Supp Fig 7A), supporting the *Myc*- transcriptional target profile identified in this cluster.

Upon characterizing the phenotypic plasticity of the aforementioned tumor states, we sought to elucidate whether these states represent continuous steps along evolutionary trajectories in K, KP+/−, and KP tumors. To do so, we applied trajectory inference algorithms^45,46^ (Methods) to K, KP+/−, and KP malignant cells (Fig 4E-G), designating the AT2-like state as the cell-of-origin in these tumors^47,48^. In K tumors, the predominant path of tumor progression was the lung injury-induced trajectory (Fig 4E; Supp Fig 7B). This trajectory was characterized by a sharp transition in pseudotime, in which the AT2-like tumor state markers were abruptly downregulated, coinciding with the immediate transcriptional activation of Basal-like state markers and the emergence of markers defining the transitional *Cryab*+ AT1-like state that arises during alveolar repair (Supp Fig 7C). In KP+/− tumors, the predominant path of tumor progression was the gastric dedifferentiation trajectory (Fig 4F, Supp Fig 7B). This trajectory revealed a gradual pseudotime ordering of continuous gene expression transitions: initial downregulation of lung AT2-like markers was followed by sequential emergence of transcriptionally dysregulated tumor states (*Mdk*+ AT2-like, *Fetub*+ Gastric AT2-like, EMT) and transition through a high-cycling state before culminating in the induction of highly plastic and high-CNV tumor states (*PI3K*/*Akt*+ Ductal-like, *Fras1*+ ECM_remodeling etc) (Fig 4F; Supp Fig 7D). Lastly, in the KP tumors, trajectory analysis also predicted two distinct evolutionary trajectories: the first recapitulates the previously published path of tumor evolution in the KP dataset, from AT2-like cells to gastric and endodermal states before undergoing epithelial-to-mesenchymal transition^41^ (EMT) (Fig 4G). The second reflects an alternative path of tumor evolution involving an earlier transition from AT2-like to the early gastric state, which directly feeds into the mesenchymal-like and metastatic mesenchymal states. The early divergence in the alternative evolutionary trajectory suggests potential early dissemination of lung tumor cells in the KP model^49,50^. Notably, both evolutionary trajectories converge on the emergence of highly aggressive mesenchymal-like states that occur only in KP tumors.

### KP+/− tumor state signatures and copy-number alterations are conserved and predict clinical outcomes in human NSCLC

To conclude our efforts on characterizing the KP+/− model of lung tumor progression, we explored the relevance of the identified mouse KP+/− tumor states and CNVs in human NSCLC. We generated tumor state signatures by identifying the distinct upregulated genes that define each of the 13 annotated tumor states (Supp Table 5). Scoring KP+/− tumor state signatures in human *KRAS*-mutant LUAD bulk-RNA sequencing data from The Cancer Genome Atlas (TCGA) revealed variable representation of the 13 KP+/− tumor-state programs across patients^23,51^ (Fig 5A). Furthermore, Cox proportional hazards modeling demonstrated that LUAD tumors with high expression of the *Mdk*+ AT2- like signature had worse survival (*P* = 0.0324), whereas the high expression of the AT2- like state correlated with a more favorable outcome (*P* = 0.0123) (Methods; Fig 5B). Kaplan–Meier analysis further showed poorer survival among patients with high *Mdk+* AT2-like signature activity in both early- and advanced-stage LUAD, whereas higher AT2- like signature activity was associated with improved survival in early-stage disease (Fig 5C; Supp Fig 8A). Notably, the direct correlation of KP+/− -specific *Mdk*+ AT2-like signature activity with poor survival outcomes highlights the clinical relevance of transcriptional states revealed by *Trp53* heterozygosity, particularly given that retention of a wild-type *TP53* allele is more prevalent than complete *TP53* loss in human *KRAS*- mutant NSCLC (Supp Fig 1A).

**Fig 5:**
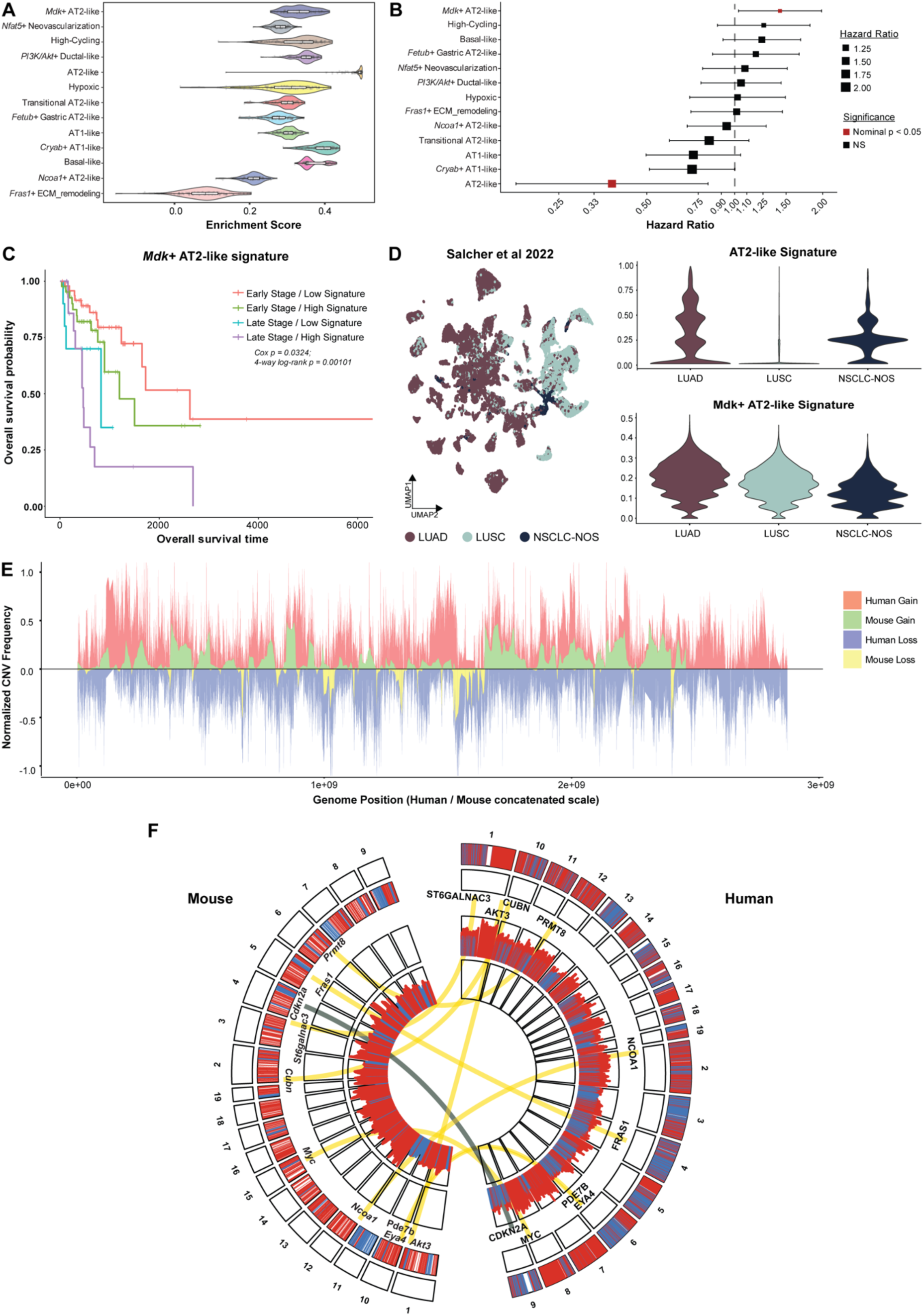
Murine KP+/− genomic profiles are conserved in human *KRAS*- mutant LUAD and associate with poor survival. **(A)** Rank-based enrichment scores of KP+/− tumor state signatures in human *KRAS*-mutant LUAD bulk-RNA sequencing data from TCGA (n = 132 patients). **(B)** Forest plot showing hazard ratios (HRs) (x-axis) and 95% confidence intervals (CIs) for overall survival associated with KP+/− tumor state signature (y-axis) expression in the TCGA KRAS-mutant LUAD cohort (n = 117 patients). Hazard ratios were estimated using Cox proportional hazards regression models, adjusted for patient age and stratified by pathological stage (early vs. late). Squares indicate hazard ratios, horizontal lines represent 95% confidence intervals, and square size is proportional to the magnitude of the hazard ratio. The dashed vertical line denotes HR = 1 (no association with survival). Red squares indicate signatures with nominal *P* < 0.05, whereas black squares indicate non-significant (NS) associations. **(C)** Kaplan–Meier overall survival curves for TCGA KRAS-mutant LUAD patients stratified by pathological stage (early *vs.* late) and *Mdk*+ AT2-like signature expression (high *vs.* low, dichotomized at the cohort median), generating four patient groups: Early Stage/Low Signature, Early Stage/High Signature, Late Stage/Low Signature, and Late Stage/High Signature. Overall survival differences among the four groups were assessed using a 4-way log-rank test. The reported Cox P-value was obtained from analyses in (B). **(D)** Left: UMAP visualization of tumor subtypes (LUAD – lung adenocarcinoma; LUSC – lung squamous cell carcinoma) in the human NSCLC scRNA-sequencing atlas from *Sachler et al*. Right: Violin plots indicate the expression of AT2-like and *Mdk*+ AT2-like gene signatures for tumor subtypes from (D). **(E)** Comparative genome-wide CNV frequency profiles in human *KRAS*-mutant LUAD (TCGA) and KP+/− mouse tumors. Whole-genome sequencing data from human *KRAS*-mutant patients (n = 132 patients) were compared to the inferred copy number alterations profile from mouse KP+/− tumor states. Gains are shown above zero and losses below zero, with human events in red (gain) and blue (loss), and mouse events in green (gain) and yellow (loss). Frequencies were normalized to a common scale and plotted across concatenated genome coordinates; dashed lines indicate chromosome boundaries. **(F)** Comparative circos plot of conserved copy number alterations between human *KRAS*-mutant LUAD and the KP+/− mouse model. The outer heatmap tracks represent genome-wide copy number alterations across mouse (left) and human (left) genomes, with red indicating copy number gains and blue indicating copy number losses. The inner histogram tracks depict the corresponding magnitude of copy number alterations across each genome. Genes that define the KP+/− mouse tumor states with high copy number burden are labeled at their genomic locations. Links connect orthologous loci between species: yellow ribbons indicate conserved copy number gains in both human and mouse, whereas the gray ribbon highlights a discordant event at the *CDKN2A* locus, characterized by copy number loss in human *KRAS*-mutant LUAD and neutrality in the KP+/− mouse tumors.

To further evaluate the expression of mouse KP+/− tumor state signatures, we used an integrated human NSCLC scRNA-seq dataset comprising 42 patients^52^. UCell-based gene signature scoring identified restricted expression of the AT2-like signature in LUAD, whereas the *Mdk*+ AT2-like signature is expressed across both LUAD and LUSC subtypes, suggesting that the *Mdk*+ AT2-like state is present broadly across NSCLC patients (Fig 5D). In addition, heterogeneity in the expression of KP+/− tumor signatures was evident in annotated tumor states in the human NSCLC data. (Supp Fig 8B).

Finally, to validate the mouse-human cross-species conservation of CNVs, we compared genome-wide CNV landscapes generated from whole-genome sequencing (WGS) data of TCGA human *KRAS*-mutant LUAD tumors (Supp Fig 8C) to the inferred mouse KP+/− tumor states^23,51^ (Supp Fig 8D). Although the overall frequency of CNVs was greater in human tumors, both mouse and human malignant cells exhibited recurrent gains and losses across orthologous genomic regions (Fig 5E). *KRAS* was a conserved orthologous gain, whereas frequent *KRAS* co-mutations in LUAD: *KEAP1*, *STK11*, and *TP53* were lost across mice and humans^11^ (Supp Fig 8E). Extending this analysis revealed that genes defining the KP+/− high-CNV tumor states (*FRAS1*, *NCOA1*, *AKT3*, *CUBN*, *PRMT8*, *EYA4*, *PDE7B*, *ST6GALNAC3*) and KP+/− *Mdk*+ AT2-like state (MYC) were recurrently gained in both mouse and human tumors, demonstrating that the genomic programs underlying the high-CNV states unique to the KP+/− model are broadly conserved across species (Fig 5F). These concordant alterations suggest that several genomic features associated with KP+/− tumor states are also represented in human *KRAS*-mutant LUAD. In contrast, the *CDKN2A* locus exhibited a species-specific discordant pattern, with recurrent loss in human LUAD but relative gain in KP+/− mouse tumors. Together, these analyses demonstrate cross-species concordance of selected transcriptional and copy-number features identified in the KP+/− model and support its clinical relevance in human KRAS-mutant NSCLC.

## Discussion

In this study, we have identified unique tumor cell states in a lung tumor model driven by oncogenic *Kras* and *Trp53* heterozygosity. The *PI3K/Akt*+ Ductal-like and *Mdk*+ AT2-like states are characterized by activation of PI3K/Akt signaling and Myc pathway activity, respectively. These signaling pathways have been implicated in driving resistance to KRAS-G12C inhibitors in LUAD patients^53,54^. Therefore, our model underscores the possibility that malignant cells driving resistance to KRAS-targeted therapies may already preexist in tumors and supports the notion of intrinsic versus acquired resistance in LUAD patients. Another unique KP+/− state: *Nfat5*+ neovascularization potentially promotes angiogenesis and capillary support by local endothelial cells to meet the vascular demands and advanced progression of KP+/− tumors^55,56^. Additionally, the tumor states shared between K and KP+/− tumors, which expand in proportion in the latter, exhibit features of aggressive tumor progression. The Basal-like tumor state was characterized by dysregulated metabolic programs, a hallmark of Kras-driven tumors^57,58^, whereas the Hypoxic cluster most closely resembled the high-plasticity cell state (HPCS) in terms of marker expression (*Cldn4*, *Msln*, *Procr*), signature activity in human MSLN+ LUAD tumor cells (Supp Fig 8B), and has been previously identified in progressing KP murine lung tumors^21,59^. Therefore, *Trp53* heterozygosity in combination with an oncogenic *Kras* mutation increases the number of cell states leading to tumor heterogeneity and potentially driving increased resistance to targeted therapies.

The genomic instability detected in KP+/− mice might result from the increased tumor latency in these mice (12-18 months), compared with only 6 months in KP mice^17^. Over time, KP+/− malignant cells acquired copy-number variations, giving rise to some of the unique tumor states. Our K and KP+/− models contain broad chromosomal instability, while in our analysis of KP tumors, we observed only focal amplifications and losses. The broad chromosomal changes in KP+/− mice, which also occur in human LUAD^60–62^, may represent an unexploited opportunity to examine the immunopeptidomic landscape and identify unique antigens arising in these malignant cells for therapeutic targeting^63^.

In our K–KP+/− integrated tumor analysis, EMT and metastatic events were absent, unlike those of KP lung tumors^41^. This suggests that lung tumors might need to lose wild type *Trp53 in vivo* for EMT and metastasis to occur. This suggestion is supported by a recent study examining Kras-mutant *Trp53* heterozygous pancreatic tumors, which show that loss of wild-type *Trp53* is necessary for mesenchymal transdifferentiation^64^.

The basis of tumoral heterogeneity in KP mice has been attributed to epigenetic changes and altered transcription factor activity, rather than copy-number events^41,43^. Our data support this model because the three states in K and KP+/− tumors that map to KP tumors (Hypoxic, *Fetub*+ Gastric AT2-like, and Basal-like) have low copy-number burden and increased TF activity, potentially reflecting a distinct chromatin landscape compared to the high CNV-burden states. Therefore, it is possible that both genetic/copy-number changes and chromatin changes serve as two distinct mechanisms driving the intertumoral heterogeneity.

Our studies demonstrate that *Trp53* heterozygosity is not a mere “halfway point” in lung Kras-mutant tumor evolution. Instead, KP+/− tumor states are characterized by copy-number alterations and phenotypic plasticity, both of which are detectable in human NSCLC data and have important implications for tumor evolution and clinical prognosis. By coupling GEMM-based single-cell transcriptional profiling with human clinical validation, this work provides new insights into how *TP53* dosage shapes KRAS-driven lung adenocarcinoma and nominates gene signatures with prognostic potential for patient stratification.

## Methods

### Mice

All mouse experiments were performed in compliance with National Institutes of Health (NIH) guidelines and Association for Assessment and Accreditation of Laboratory Animal (AAALAC) accreditation standards for animal research and approved by The University of Texas MD Anderson Cancer Center Institutional Animal Care and Use Committee. The Kras^LSL-G12D^ mice were obtained from Dr. Jonathan M. Kurie’s lab. The p53^fl^ allele (Strain #: 008462) was obtained from the Jackson Laboratory.

Lenti-Cre-Luc2-P2A-tdTomato vector was generated by cloning the Luc2-P2A-tdTomato fragment from pCDH-EF1-Luc2-P2A-tdTomato (Addgene #72486) using EcoRI and XhoI restriction enzyme cut sites and inserted into Puro.Cre vector (Addgene #17408). Lenti-Cre-Luc2-P2A-tdTomato lentivirus was produced by transfecting Lenti-Cre-Luc2-P2A- tdTomato vector together with psPAX2 (Addgene #12260) and pCMV-VSV-G (Addgene #8454) vectors in 293T cells. The functional titer of the produced lentiviral particles was determined by analyzing the percentage of RFP positive cells using Flow cytometry after transducing 293T cells with the produced lentivirus. Sex of the mice did not significantly influence the resulting tumor transcriptome analysis (Supp Table 1). Mice were euthanized in this study only when they exhibited hyperpnea or breathing difficulty.

### Tumor Burden Quantification

Hematoxylin and Eosin staining of tumor-bearing whole-lung sections from K and KP+/− was performed as previously published. The Vectra Polaris microscope was used to image whole-lung sections at 10x magnification. Every adenoma and adenocarcinoma within each lung section was manually annotated, and the cross-sectional area of each tumor (µm²) was measured to calculate the tumor size. Tumor burden was expressed as the percentage of total lung area occupied by tumors. Tumor burden was calculated by dividing the cumulative area of all annotated tumors by the total lung area and multiplying by 100. Histopathological grading of tumors was performed by an in-house pathologist.

### Single-Cell Dissociation of tumor-bearing whole lungs

Following euthanasia, murine lungs were thoroughly perfused with phosphate-buffered saline (PBS) via the right ventricle to clear circulating blood cells. Tumor-bearing whole lungs were harvested into Gibco™ Leibovitz’s L-15 Medium (Gibco, cat. no. 21083-027) and finely minced using forceps under the dissection microscope. To obtain single-cell suspensions, minced tissue was subjected to both enzymatic digestion in L-15 medium supplemented with 2mg/mL Collagenase Type I (Worthington, cat. no. LS004197), 2mg/mL Elastase (Worthington, cat. no. LS002294), and 0.5mg/mL DNase I (Worthington, cat. no. LS002007) for 30 minutes on a 37°C water bath. The tissue was mechanically triturated midway through the digestion period (15 minutes). Enzymatic activity was quenched by adding fetal bovine serum (FBS) to a final concentration of 20%. Following quenching, samples were placed on ice and maintained at 4°C for all subsequent steps. The cell suspension was filtered through a 70 µm cell strainer (Falcon, cat. no. 352350) and centrifuged at 1500 rpm for 1 minute. To eliminate remaining erythrocytes, the pellet was resuspended in ACK red blood cell lysis buffer (Gibco, cat. no. A1049201) for 3 minutes. Cells were pelleted (5000 rpm for 1 minute), washed with L-15 medium containing 10% FBS, and passed through a 35 µm strainer cap (Falcon, cat. no. 352235). Single cells were centrifuged and resuspended in 1X PBS with 0.04% Bovine Serum Albumin (BSA).

### scRNA-sequencing of tumor-bearing whole lungs

Cell viability was assessed using 0.4% Trypan Blue staining with the Countess™ II FL automated cell counter. Cells were diluted or concentrated to the recommended range of 500–1,000 cells/µL and loaded at a volume to target a recovery of 7,000–10,000 cells per sample for single-cell RNA sequencing (scRNA-seq). Single-cell capture, barcoding, and library preparation were performed according to the 10x Genomics Chromium Next GEM Single Cell 3′ v3.1 Reagent Kits protocol (CG000204 Rev D). Ten libraries were pooled to a final concentration of 10 nM. The pooled libraries were further quantified by qPCR prior to sequencing. Sequencing was performed on an Illumina NovaSeq 6000 using an S2 100-cycle flow cell, with 28 cycles for Read 1, 8 cycles for the i7 index, and 91 cycles for Read 2. Sequencing was conducted at the ATGC Core Facility at MD Anderson. The number of reads obtained for every sequenced biological replicate is reported in Supplementary Table 1.

### Computational Analysis

#### scRNA-Seq Processing and Quality Filtering

Raw sequencing data were demultiplexed and aligned to a custom mouse genome (mm10) containing *tdTomato*, *Cre* and *Luciferase* reporter sequences using the Cell Ranger v7.0.1. Unique molecular identifier (UMI) count matrices generated by Cell Ranger were analyzed using the Seurat package (v.5.1.0) in R. Low-quality cells and cell debris with >15% of UMI counts mapping to mitochondrial genes, and <200 detected genes were excluded from downstream analyses. Cells with >7,500 detected genes were also considered potential doublets or multiplets and excluded. The upper gene-detection threshold of 7500 was selected by examining the relationship between total UMI counts (nCount_RNA) and the number of detected genes (nFeature_RNA) across samples, filtering out cells with abnormally high values for both metrics (Supplementary Table 1). Filtered UMI count matrices were normalized using NormalizeData, and highly variable genes were identified using FindVariableFeatures. Data were scaled using ScaleData, with default Seurat parameters unless otherwise indicated. Principal components Analysis (PCA) was performed, and the number of dimensions was estimated using the JackStraw and ElbowPlot. A shared nearest-neighbor graph was constructed using FindNeighbors, followed by graph-based clustering using FindClusters. Uniform manifold approximation and projection (UMAP) was performed using RunUMAP for visualization. Additional quality filtering was performed iteratively following clustering. Clusters showing discordant co-expression of markers from multiple major lineages were considered putative doublets and excluded. Lineage markers included *Ptprc* for immune cells; *Col1a1*, *Col3a1*, *Pdgfra*, and *Pdgfrb* for mesenchymal cells; *Epcam*, *Cdh1*, *Krt8*, and *Krt18* for epithelial cells; and *Plvap*, *Cdh5*, and *Icam2* for endothelial cells. Clusters lacking expression of canonical lineage markers were considered low-quality or debris populations and were also excluded. Differentially expressed genes for each cluster were identified using FindAllMarkers, and clusters whose top differentially expressed genes were predominantly mitochondrial or ribosomal genes were removed as additional low-quality populations. Following quality filtering, epithelial, immune, mesenchymal, endothelial, and proliferating cell populations were retained for downstream analyses. Cellular composition was visualized using the do_BarPlot function from the SCpubr package. Proliferating cells were classified as a cellular state rather than a distinct lineage and predominantly expressed immune-lineage markers.

#### Malignant cell identification in K and KP+/− tumor-bearing scRNA-seq data

Batch effect correction was performed using the Harmony package (v1.2.1) when integrating K and KP+/− single-cell datasets^26^. First, we subset the epithelial lineage cells from the global K and KP+/− data, cluster at resolution 1.5 using the FindClusters function, and filtered out any non-epithelial cells. To distinguish normal epithelial populations from tumor-associated epithelial cells, the .h5ad version of the normal murine lung cell atlas dataset from LungMAP^31^. The reference dataset was normalized, highly variable features were identified, the data were scaled, and PCA and graph-based clustering were performed as described above. The processed LungMAP object was then converted to an RDS object (Epithelial_LungMAP). Cell-type labels were transferred from the LungMAP reference to K and KP+/− epithelial cells using FindTransferAnchors and TransferData, with Epithelial_LungMAP as the reference and the K and KP+/− epithelial cells as the query.

As an independent annotation approach, published gene signatures^28,29^ for normal lung epithelial cell types (basal, club, ciliated, AT1, and AT2 cells) were scored across all K and KP+/− epithelial cells using the UCell package (v2.10.1). Cluster-level mean UCell scores, transferred LungMAP annotations, and expression of canonical epithelial markers were jointly used to annotate normal epithelial populations and distinguish them from tumor-associated epithelial states. Finally, DotPlot was used to visualize the percentage of cells expressing, and the average expression of, canonical epithelial cell-type markers across annotated normal epithelial populations and tumor states. The number of normal and malignant epithelial cells for every sample is reported in Supplementary Table 1.

### K and KP+/− tumor state analyses

#### STRING analyses

Malignant epithelial cells were subset from the K and KP+/− epithelial datasets and reintegrated using Harmony. The integrated malignant-cell object was joined using the JoinLayers function, followed by differential expression analysis between KP+/− and K malignant cells using FindMarkers with the MAST test. Following differential expression analysis, predicted genes and ribosomal and mitochondrial genes were excluded from downstream pathway analysis to reduce technical and housekeeping-associated signals. Genes with a log2 fold change >1.5 and an adjusted P value <0.05 were selected as genes upregulated in KP+/− relative to K malignant cells and subjected to pathway and protein–protein interaction network analysis using Metascape/STRING^30^ (Supp Table 2). All genes detected across K and KP+/− malignant cells were used as the background gene set for enrichment analysis. Network results were visualized using Cytoscape. Nodes represent enriched pathways, with node color indicating network membership and node size representing the enrichment score. Related pathways were grouped into distinct networks.

#### Gene expression and Gene set enrichment analysis (GSEA)

Then, we reclustered the subset K and KP+/− malignant cells at a resolution of 0.5, yielding 13 discrete tumor-state clusters. Clusters were annotated using canonical lineage markers and published gene signatures representing neural, liver, intestinal, AT1, and AT2, as well as cell-cycle and hypoxia programs^31–39^. Differential expression analysis was performed for each cluster using the FindAllMarkers function with the MAST test while controlling for genotype as a latent variable. Mitochondrial, ribosomal, and predicted genes were excluded, and genes were ranked by average log2 fold change for preranked GSEA using the fgsea (v1.32.0) package^40^. Gene sets were obtained from the Molecular Signatures Database (MSigDB) using the msigdbr (v7.5.1) package, including the Hallmark (H), Canonical Pathways (C2; KEGG, WikiPathways, and BioCarta), Gene Ontology (C5; Biological Process, Molecular Function, and Cellular Component), Cancer Gene Neighborhood and Cancer Modules (C4; CGN and CM), and Cell Type Signature (C8) collections. Positive enrichment analysis was performed (scoreType = “pos”), and significantly enriched pathways were identified based on adjusted *P* values (FDR < 0.05) (Supp Table 3). A list of differentially upregulated and downregulated genes in KP+/− Basal-like, Hypoxic, and *Fras1*+ ECM_remodeling clusters compared to the K counterpart can be found in Supplementary Table 4. Pathways that are trending or statistically enriched in the expanding KP+/− tumor states compared to the K counterpart are also reported similarly (Supp Table 4).

#### Bidirectional label transfer between K - KP+/− and KP tumor states

The published KP scRNA-seq dataset were obtained as a preprocessed .h5ad object^41^ and reprocessed using normalization, scaling, PCA, and UMAP. Reference UMAP models were generated for both the KP dataset and the integrated K–KP+/− dataset using RunUMAP. Bidirectional label transfer was then performed using FindTransferAnchors, TransferData, and MapQuery. First, KP was used as the reference and K–KP+/− as the query, followed by the reciprocal analysis with K–KP+/− as the reference and KP as the query. Correspondence between tumor-state annotations was visualized using alluvial plots generated with the ggalluvial package, with flow widths proportional to the number of mapped malignant cells. K–KP+/− tumor-state signatures were defined using 4–20 genes selectively expressed in each of the 13 tumor states (Supp Table 5). KP tumor-state signatures were defined using the top 20 highly expressed genes reported for each cluster in the published KP dataset.

#### Copy number inference

Large-scale copy number variations (CNV) were inferred from single-cell transcriptomes using the inferCNV (v1.22.0) package. Age-matched wild-type CD4+ and CD8+ T-cells (18–21 months) from the Tabula Senis atlas were used as the diploid reference population for the inference^42^. Raw expression matrices from K, KP+/−, and KP malignant cells, together with tumor-state annotations and genomic gene coordinates, were used to construct the inferCNV object. Copy number inference was then performed, according to the standard inferCNV workflow (cutoff = 0.1, window_length = 101, cluster_by_groups = TRUE, denoise = TRUE, HMM = TRUE, analysis_mode = “samples”), generating denoised chromosome-scale copy number profiles and hidden Markov model-based CNV state predictions for each tumor cell.

To quantify genome-wide copy-number burden, an absolute CNV score was calculated for every malignant cell by summing the absolute deviation of inferCNV expression values from the inferred diploid reference baseline (|CNV value − 1|) across all genes. Absolute CNV scores were added as a metadata column and visualized on the UMAP to compare copy-number burden across annotated tumor states in K, KP+/−, and KP genotypes. *Kras* and *Trp53* inferCNV values for all malignant cells were extracted from the resultant inferCNV expression matrix and added to the Seurat metadata as well. Mean (average) inferCNV-derived copy-number values for *Kras* and *Trp53* were computed for each annotated tumor state and compared across K and KP+/− (*Trp53*) alongside KP genotypes to assess locus-specific copy-number alterations. Decrease in inferCNV-derived *Trp53* copy-number values in KP+/− compared to K annotated tumor states was assessed using two-sided Wilcoxon rank-sum tests. Mean inferCNV values ± standard error is displayed for visualization.

#### Inference of tumor plasticity in scRNA-seq data

Cellular differentiation potential was inferred using the CytoTRACE package^44^ (v0.3.1) from raw expression matrices of malignant cells from integrated K - KP+/− and KP datasets. CytoTRACE scores were calculated for each malignant cell using default parameters and incorporated into the corresponding Seurat metadata. Differences in CytoTRACE scores between KP+/− and K tumor states were assessed using two-sided Wilcoxon rank-sum tests.

#### Transcription factor activity analysis

Transcription factor (TF) activities were inferred from the integrated K - KP+/− single-cell transcriptomic dataset using the decoupleR (v2.12.0) package with the CollecTRI regulatory network. Human CollecTRI regulons were mapped to mouse orthologs, and per-cell TF activity scores were estimated using the weighted mean (run_wmean) method with 100 permutations and a minimum regulon size of five target genes. Mean TF activity was calculated across the 13 annotated tumor states, and highly variable TFs were visualized by hierarchical clustering of z-scored activity values. Inferred *Myc* activity and *Myc* gene expression were additionally visualized across malignant-cell states.

#### Inference of tumor evolutionary trajectories in scRNA-seq data

Pseudotime trajectories were inferred separately for K, KP+/−, and KP malignant cells using the Monocle3 package^45^ (v1.4.27). Seurat objects were converted to Monocle objects, followed by cell clustering, principal graph learning, and pseudotime ordering. The AT2-like tumor state was selected as the trajectory root for each genotype, and inferred trajectories were visualized on the corresponding UMAP embeddings.

Probabilistic trajectory analysis was performed on the integrated K - KP+/− tumor dataset using the Palantir package^46^. Normalized RNA expression and cell metadata were transferred from Seurat to an AnnData object, and the 2,000 most variable genes were used for principal component and diffusion-map analyses. An AT2-like cell was selected as the trajectory root, and Palantir was used to estimate pseudotime, differentiation entropy, terminal states, and branch probabilities. Two major trajectories were annotated based on tumor-state identities and marker-expression patterns. Branch-specific gene-expression dynamics were estimated using Markov Affinity-based Graph Imputation of Cells (MAGIC)- smoothed expression values and visualized as gene-wise z-scored trends across pseudotime.

#### Human clinical data analyses

First, overall survival data were obtained from cBioPortal across all available NSCLC studies^24,25^. Patients with KRAS-mutant tumors were stratified according to TP53 copy-number status as wild-type, heterozygous loss (HETLOSS), or homozygous deletion (HOMDEL), and overall survival was compared among groups.

For TCGA-LUAD analyses, somatic mutation data, STAR gene expression files, and clinical annotations were obtained from the NCI Genomic Data Commons via TCGAbiolinks^23,51^. Patients with non-silent KRAS mutations and matched primary tumor RNA-seq data were retained. Gene expression values were represented as log_2_(TPM + 1), with duplicate gene symbols collapsed to the mean expression.

Activities of murine-derived KP+/− tumor-state signatures were quantified in each tumor using the rank-based singscore method (Supp Table 5). Cox proportional hazards models were used to evaluate associations between continuous, z-score-standardized signature activities and overall survival, adjusting for age at diagnosis and pathological stage. Hazard ratios represent the change in mortality risk associated with a one-standard-deviation increase in signature activity. For visualization, signatures were dichotomized at the cohort median, and survival distributions were compared using Kaplan–Meier analysis and log-rank tests. Multiple-testing correction was performed using the Benjamini–Hochberg false discovery rate method. Survival analyses were performed using the survival and survminer packages.

#### Human scRNA-seq data comparisons

The processed human NSCLC scRNA-seq atlas from Salcher et al^52^ was obtained as an .h5ad object and converted to a Seurat object. Analyses were restricted to malignant cells annotated in the original study. Human Ensembl identifiers were converted to gene symbols, and murine KP+/− tumor-state signatures were mapped to their human orthologs before scoring with UCell. Original tumor-state and histological annotations were retained for downstream analyses. Signature activities were compared across LUAD, LUSC, and mixed NSCLC tumors. Selected KP+/− tumor-state signatures were visualized using violin plots, and mean z-score-standardized activities of all 13 signatures across published tumor states were summarized using hierarchically clustered heatmaps.

#### Comparative copy number alteration analyses

Whole-genome sequencing (WGS) copy number segmentation data and somatic MAF for the TCGA-LUAD cohort were obtained from the NCI GDC^23,51^. Primary lung adenocarcinoma tumors harboring non-silent *KRAS* mutations (n = 132) were retained for the analyses. Genome-wide copy number alteration (CNA) frequencies were calculated in 1-Mb genomic bins using segmented copy number values. Copy number gains and losses were defined using segment mean thresholds of > 0.2 and < −0.2, respectively, and the proportion of tumors exhibiting gains or losses within each genomic bin was calculated to generate genome-wide CNA frequency profiles.

Murine CNA profiles were inferred from malignant tumor cells using inferCNV, as previously described. Gene-level inferCNV values were mapped to GRCm39 genomic coordinates and grouped into 1-Mb intervals. Copy-number gains and losses were defined using inferCNV thresholds of >1.1 and <0.9, respectively, and their frequencies across cells or samples were calculated for each interval and visualized across mouse chromosomes 1–19.

For gene-level cross-species comparisons, human gene coordinates were obtained from EnsDb.Hsapiens.v86, and TCGA copy-number segments were assigned to overlapping genes using GenomicRanges. When multiple segments from the same tumor overlapped a gene, segment values were averaged. Gene-level gain and loss frequencies were then calculated across the KRAS-mutant LUAD cohort. Corresponding mouse gene-level CNA frequencies were calculated from inferCNV estimates using the same gain and loss thresholds described above.

One-to-one human–mouse orthologs were obtained from Ensembl BioMart and matched to gene-level CNA frequencies in each species. Net CNA frequency was calculated as the difference between gain and loss frequencies. Cross-species CNA patterns were visualized using circos plots generated with the circlize package, with separate tracks displaying gain and loss frequencies in human KRAS-mutant LUAD and KP+/− mouse tumors. Canonical NSCLC-associated genes, including *TP53, STK11, and KEAP1*, as well as genes associated with the copy-number-high KP+/− tumor states, were highlighted to assess concordant recurrent CNA patterns at orthologous loci.

#### Quantification and statistical analysis

Statistical details of experiments can be found in the figure legends and main text above. When appropriate, multiple hypothesis testing correction was employed using the Benjamini-Hochberg procedure at a false discovery rate (FDR) < 0.05.

## Data and code availability

Raw scRNA-seq data and filtered feature–barcode matrices were deposited to the Gene Expression Omnibus under accession number GSE343294. Processed single-cell data and scripts used for downstream analyses and figure generation will be publicly available through Zenodo upon publication.

---

**Supp Fig 1:**
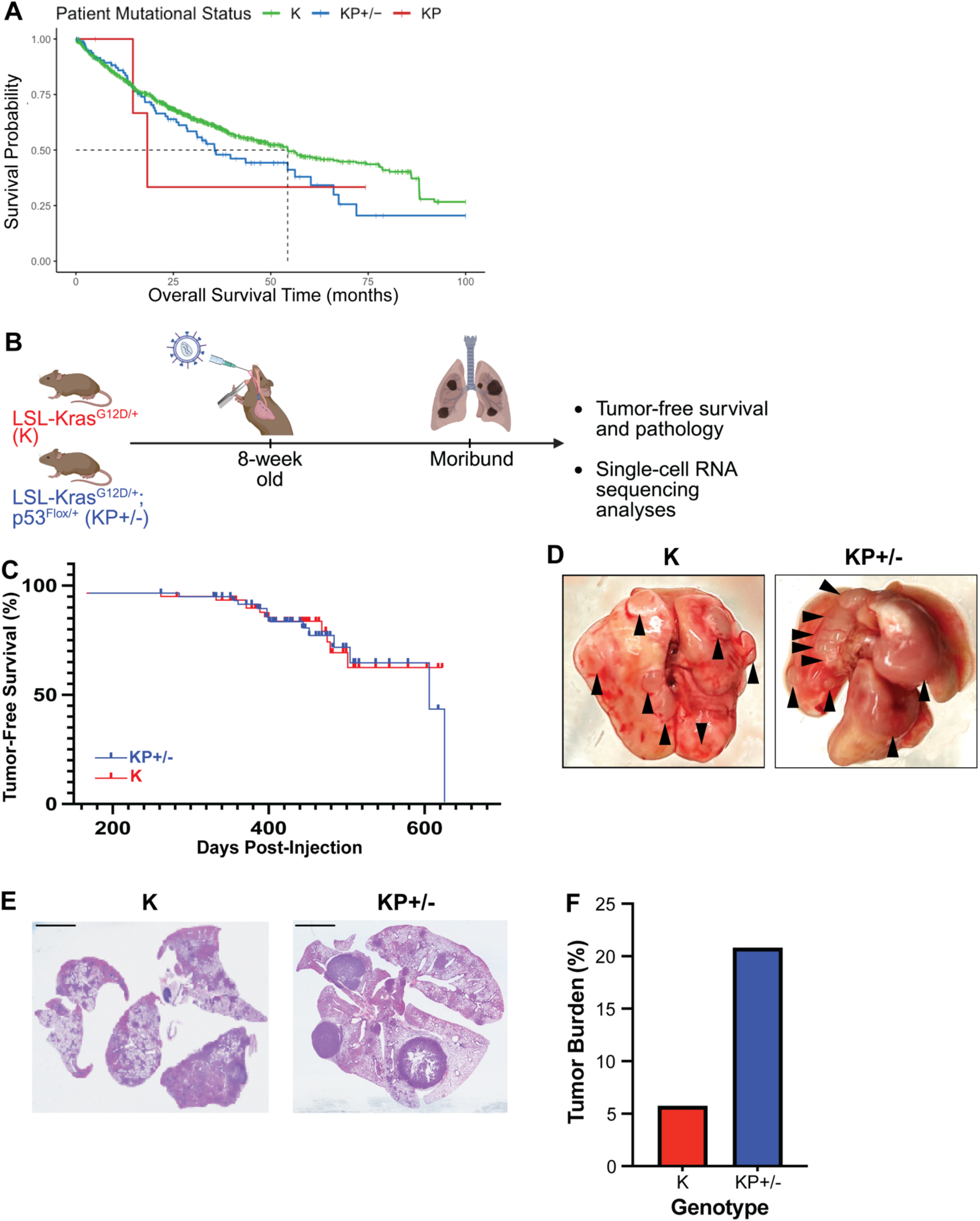
p53 dosage influences tumor survival and burden in Kras-initiated lung cancers. **(A)** Survival probabilities of non-small cell lung cancer (NSCLC) patients with *KRAS* mutations and either wild-type p53 (K [green], n = 1550 patients), single-allele loss of p53 (KP+/− [blue], n = 100 patients), and complete loss of p53 (KP [red], n = 4 patients). **(B)** Schematic diagram of the experimental design for this lung cancer study utilizing Kras^G12D/+^ (K, n=14) and Kras^G12D/+^; p53^Flox/+^ (KP+/−, n=14) mice. K and KP+/− mice were intratracheally injected with Cre-encapsulated lentivirus at 8 weeks of age. Moribund mice were sacrificed (due to breathing difficulty), and tumor-bearing lungs were collected for tumor-free survival and pathology analyses, and single-cell RNA (scRNA) sequencing. **(C)** Tumor-free survival times of K (red) and KP+/− (blue) mice post-tumor induction. **(D)** Representative images reflected the gross morphology of a tumor-bearing lung from moribund K and KP+/− mice. Black arrowheads indicate lung tumors. **(E)** Representative hematoxylin and eosin (H&E) staining of tumor-bearing lungs from moribund K and KP+/− mice. Images were taken at 10x magnification. Scale bar, 100 microns. **(F)** Quantification of tumor burden accounted by every tumor nodule in K and KP+/− tumor-bearing lungs from (E), as a proportion of the area of the total lung.

**Supp Fig 2:**
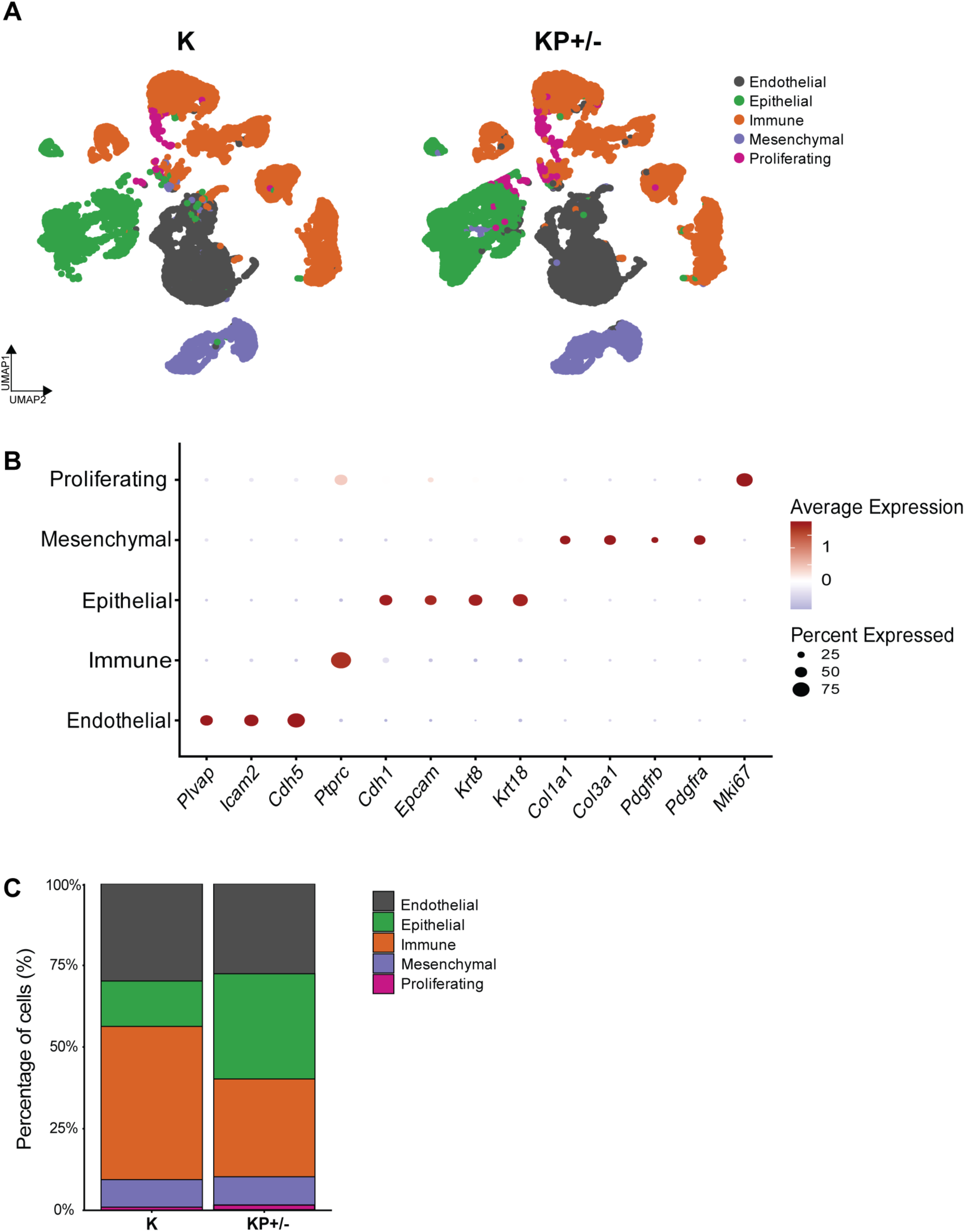
Identification of cellular lineages in scRNA-sequencing data from K and KP+/− tumor-bearing lungs. **(A)** UMAP visualization of the major clusters identified in scRNA-sequencing analyses of K and KP+/− tumor-bearing lungs (n = 3 mouse lungs per genotype - K = 21361 cells; KP+/− = 22819 cells). The data comprised the epithelial (green), endothelial (charcoal), immune (orange), mesenchymal (purple), and proliferating (magenta) lineages. **(B)** Dot plot indicate the markers (x-axis) utilized to define the major cell lineages (y-axis). The size of the dots reflected the percentage of cells expressing the marker gene. The color of the dots indicated the relative expression of the marker gene. **(C)** Stacked bar plot indicated the percentage of cells comprising the major cell lineages in K and KP+/− tumor-bearing lungs.

**Supp Fig 3:**
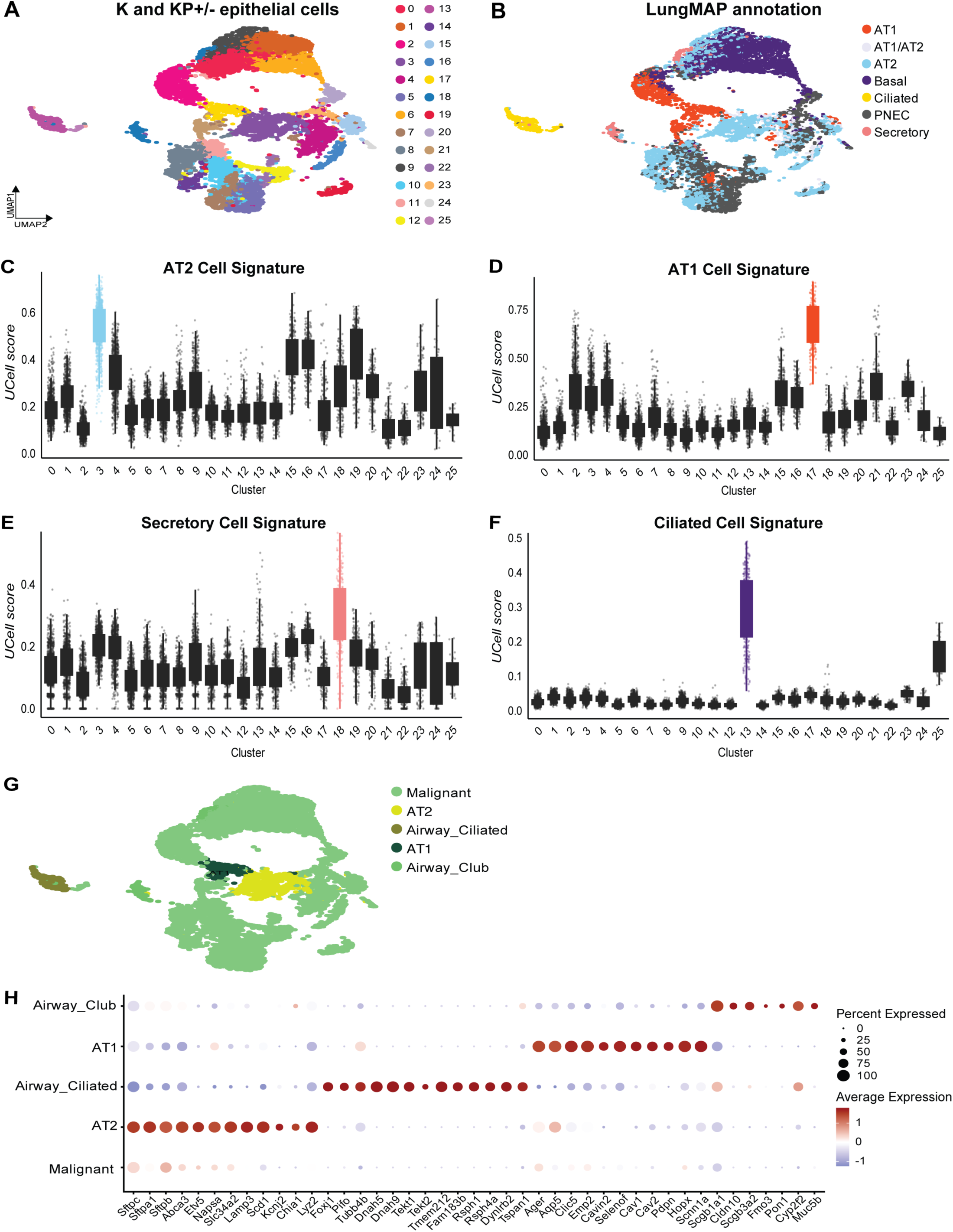
Differentiation of malignant from normal epithelial cells in tumor-bearing K and KP+/− lungs. **(A)** UMAP visualization of K and KP+/− epithelial clusters (K = 2824 cells; KP+/− = 7211 cells) post sub-clustering from the K and KP+/− epithelial compartments (from Supp Fig 2). **(B)** UMAP representation of cell type labels transferred from the epithelial compartment of LungMAP, a normal lung cell reference atlas, onto the K and KP+/− epithelial clusters. **(C-F)** Boxplots reflect the mean UCell score of cells in every epithelial cluster (from (A)) for expression of published single-cell signatures of (C) normal AT2, (D) AT1, (E) club, and (F) ciliated cells. **(G)** Annotation of malignant and normal epithelial cell types in the K and KP+/− epithelial compartments**. (H)** Dot plot of *bona fide* gene markers identifies normal AT1, AT2, airway ciliated, airway club, and malignant cell types.

**Supp Fig 4:**
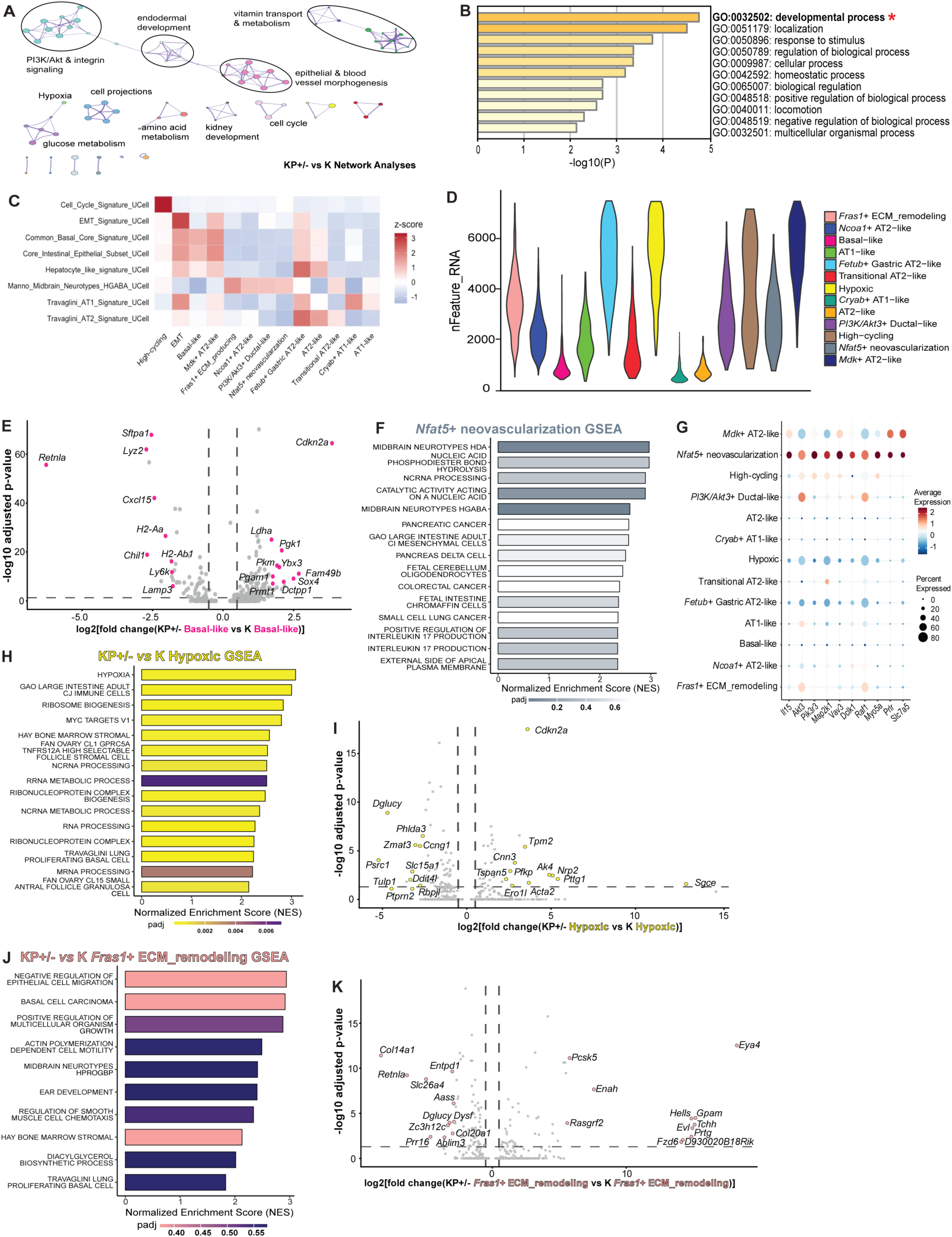
Transcriptomic characterization and pathway enrichment analyses of tumor states in K and KP+/− malignant cells. **(A)** STRING pathway interaction network analysis reveals developmental and metabolic differences, and perturbations in *PI3K/Akt* receptor-growth factor signaling in KP+/− malignant cells compared with K counterparts. **(B)** Bar chart of the top-scoring GO biological processes from STRING; the GO process with the greatest log fold change is indicated with a red asterisk. **(C)** Heatmap of Z-scores of published gene signatures’ expression in each of the thirteen annotated tumor states (from B). **(D)** Violin plot of feature counts (nFeature_RNA) across all thirteen clusters. **(E)** Volcano plot shows differentially expressed genes (DEGs) in the KP+/− basal-like tumor state compared with the K basal-like state. The top ten most up- and downregulated genes are colored. **(F)** Barplot reflects the Normalized Enrichment Scores (NES) from Gene Set Enrichment Analyses (GSEA) for functional pathways enriched in the *Nfat5*+ neovascularization tumor state. **(G)** Dot plot of leading genes overrepresented in the functional pathways in the *Nfat5*+ neovascularization tumor state (from Supp Fig 1F). **(H)** The bar plot shows NES from GSEA for functional pathways enriched in the Hypoxic state in KP+/− tumors compared to K tumors. **(I)** Volcano plot identifies DEGs in the KP+/− Hypoxic tumor state compared to the K counterpart (from Supp Fig 1H). **(J)** The bar plot shows NES from GSEA for functional pathways enriched in the *Fras1*+ ECM_remodeling state in KP+/− tumors compared to K tumors. **(K)** Volcano plot identifies DEGs in the KP+/− *Fras1*+ ECM_remodeling state compared to the K counterpart (from Supp Fig 1J).

**Supp Fig 5:**
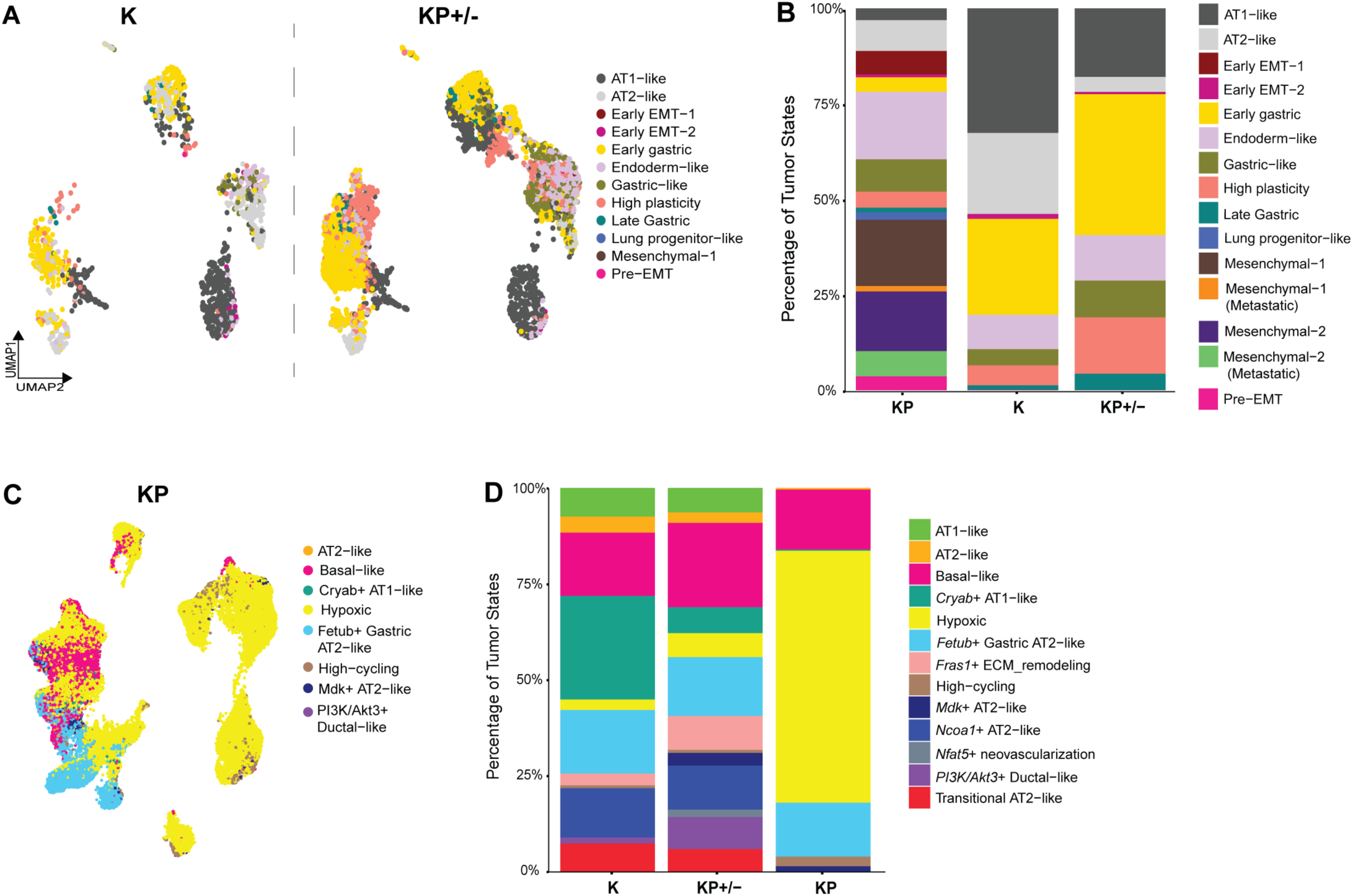
State-space validation of genotype-specific tumor states through reciprocal label transfer. **(A)** UMAP visualization of malignant cells from K and KP+/− tumors, colored according to their mapped labels from the KP (*Yang et al*) reference. **(B)** Stacked barplot indicated the percentage composition of KP tumor states across K, KP+/− and KP malignant cells. **(C)** UMAP visualization revealing malignant cells from KP tumors, colored according to their mapped labels from the K and KP+/− references. **(D)** Stacked barplot indicated the percentage composition of K and KP+/− tumor states in KP malignant cells.

**Supp Fig 6:**
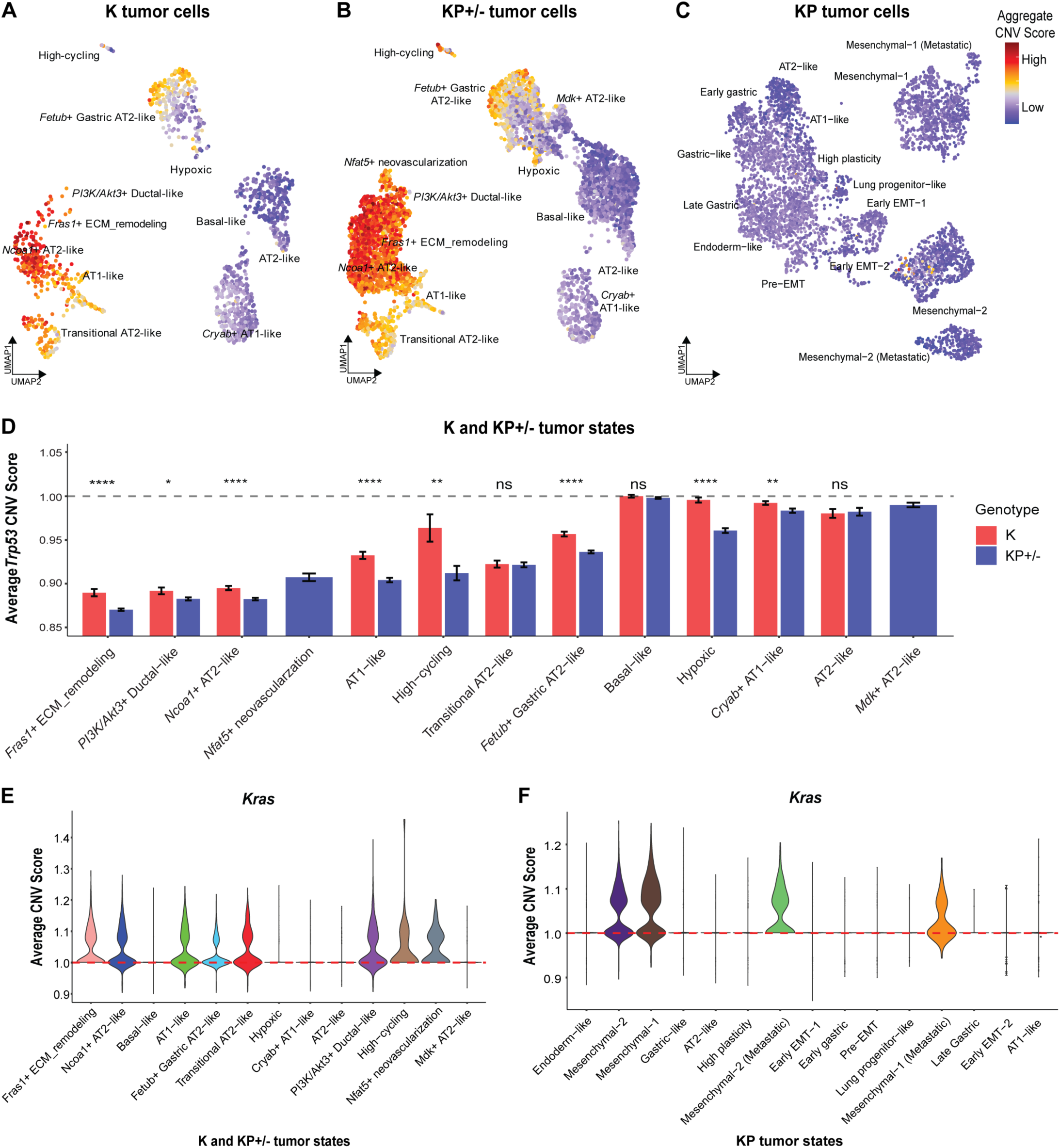
Genomic instability and CNV dynamics in *Kras*-mutant lung tumors with *Trp53* loss. **(A-C)** UMAP visualizations of (A) K, (B) KP+/−, and (C) KP malignant cells, colored by their aggregate or total Copy Number Variation (CNV) burden score (in absolute values). **(D)** Bar plots of mean *Trp53* CNV scores across the thirteen tumor states for K and KP+/− genotypes, ranked from lowest to highest score. Asterisks (*) denote a statistically significant decrease in the *Trp53* copy number in KP+/− relative to the K tumor states. **(E-F)** Violin plots reflect the inferred CNV scores for the *Kras* oncogene across K - KP+/− tumor states and KP tumor states. The red dashed line at *1.0* indicates copy number neutrality. Statistical significance was calculated using a two-tailed Wilcoxon rank-sum test, where * < 0.05, ** < 0.001, *** <0.0005, **** < 0.00001.

**Supp Fig 7:**
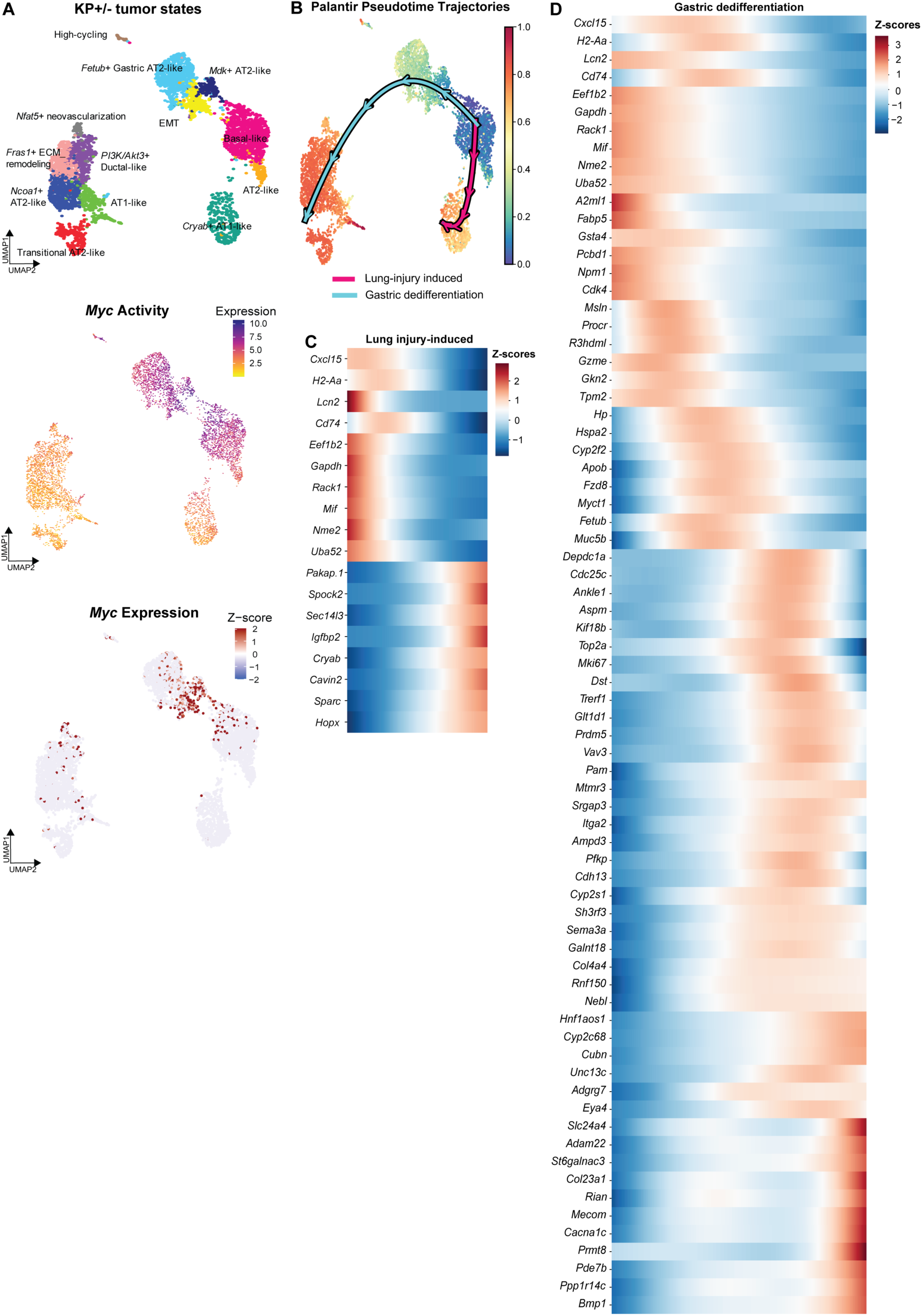
Distinct transcription factor profile and trajectory branches exist in KP+/− tumors. **(A**) UMAP visualizations of inferred transcription factor activity and expression of *Myc* across the 13 annotated tumor states. **(B)** Palantir-based pseudotime analyses of K and KP+/− tumor states. The arrows indicate the two major branches of evolution in K and KP+/− tumors, with AT2-like cells as the cells of origin. The blue-green shades reflect tumor clusters that arise earlier in tumor evolution, and the red-orange shades reflect newer states that arise later in this process. The magenta and teal arrows indicated two distinct evolutionary trajectories. **(C)** Z-scored expression of genes in tumor states ordered along the lung injury-induced trajectory from the AT2-like state. **(D)** Z-scored expression of genes in tumor states ordered along the gastric dedifferentiation trajectory from the AT2-like state.

**Supp Fig 8:**
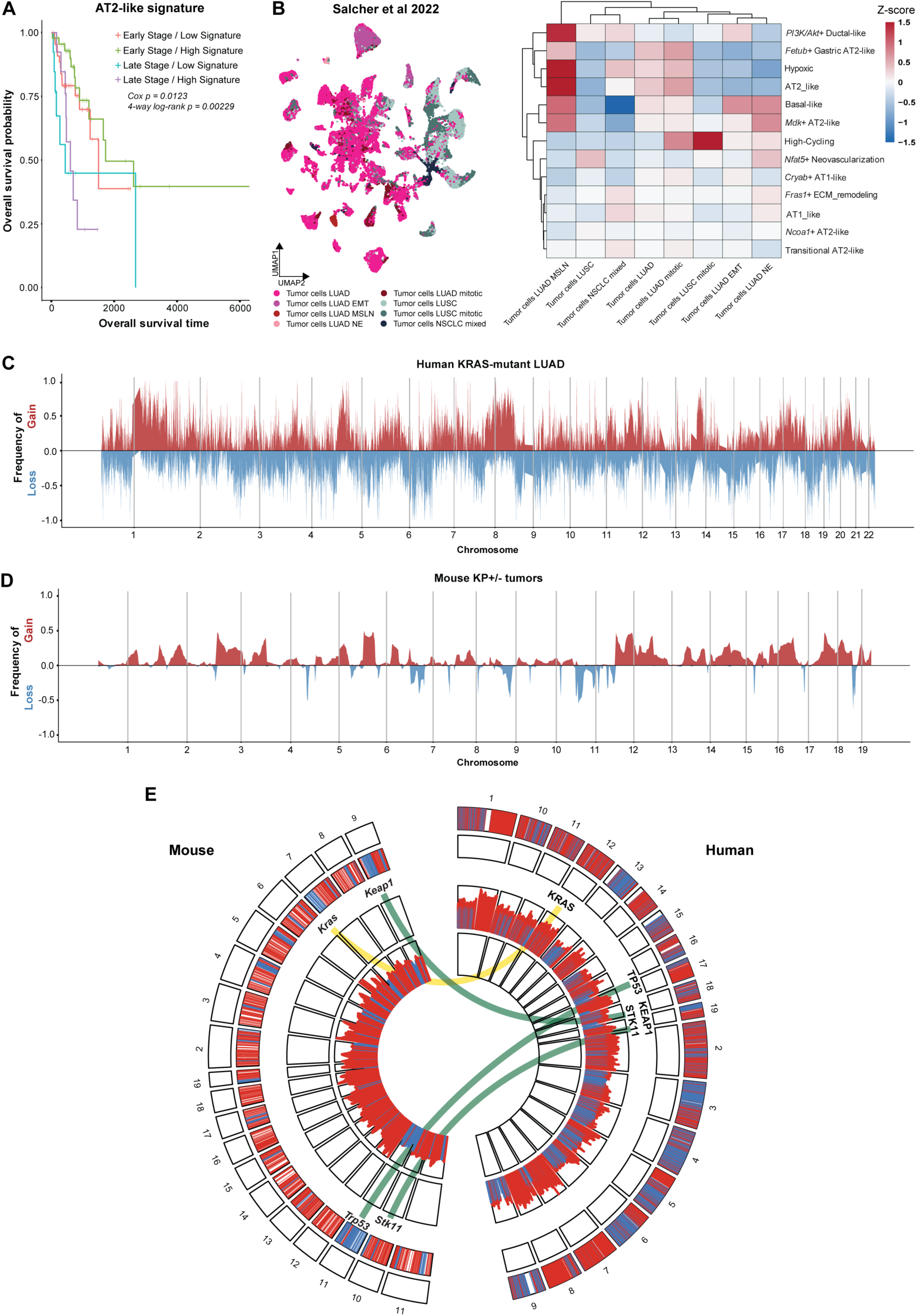
Mouse-human comparisons of KP+/− signatures and Kras co-mutation status in NSCLC genomic datasets. **(A)** Kaplan–Meier analysis of overall survival in *KRAS*-mutant human LUAD patients and AT2-like signature expression (low vs. high). Signature expression was dichotomized at the cohort median, and survival was analyzed using a Cox proportional hazards model adjusted for age and stratified by pathological stage (early vs late). Cox and four-group log-rank p-values are as indicated. **(B)** Left: UMAP projection of tumor epithelial cell states from the Salcher *et al* NSCLC single-cell transcriptomic atlas, colored by published tumor cell state annotations. Right: Heatmap of the thirteen mouse KP+/− signature scores across published tumor state cell states. Color scale indicates row-wise normalized signature scores (Z-scores). **(C)** Genome-wide CNA frequency profile of human *KRAS*-mutant LUAD patients (n = 132 patients) from TCGA whole-genome sequencing data. Copy number gains (red) are above the baseline (0) and copy number losses (blue) below the baseline across mouse chromosomes (1–19). Frequencies represent the proportion of tumor cells exhibiting gains or losses within 1-Mb genomic bins. **(D)** Genome-wide copy number alteration (CNA) frequency profiles of KP+/− mouse tumors from inferCNV analyses. Copy number gains and losses are plotted identically to Supp Fig 8C. **(E)** Comparative circos plot of genome-wide CNAs in *KRAS*-mutant human LUAD and KP+/− mouse tumors. Outer heatmap tracks reflect copy number gains (red) and losses (blue), with inner histograms depicting CNA magnitude across the genome. The plot shows KRAS and KRAS co-mutations in LUAD (TP53, STK11, KEAP1), with the yellow ribbon indicating conserved copy number gains between species and the green ribbons indicating conserved copy number loss at the designated loci.

